# Pyridoxal Kinase Inhibition by Artemisinins Downregulates Inhibitory Neurotransmission

**DOI:** 10.1101/2020.04.05.026310

**Authors:** Vikram Babu Kasaragod, Anabel Pacios-Michelena, Natascha Schaefer, Fang Zheng, Nicole Bader, Christian Alzheimer, Carmen Villmann, Hermann Schindelin

**Author notes:** These authors contributed equally to this work (Vikram Babu Kasaragod, Anabel Pacios-Michelena). Neurobiology Division, MRC Laboratory of Molecular Biology, Francis Crick Avenue, Cambridge Biomedical Campus, Cambridge, CB2 0QH, United Kingdom. **Correspondence** (V.B.K) and (H.S).

## Abstract

The anti-malarial artemisinins have also been implicated in the regulation of various other cellular pathways. Despite their widespread application, the cellular specificities and molecular mechanisms of target recognition by artemisinins remain poorly characterized. We recently demonstrated how these drugs modulate inhibitory postsynaptic signaling by direct binding to the scaffolding protein gephyrin. Here, we report the crystal structure of the central metabolic enzyme pyridoxal kinase (PDXK), which catalyzes the production of the active form of vitamin-B6 (also known as pyridoxal 5’-phosphate, PLP), in complex with artesunate at 2.4-Å resolution. Partially overlapping binding of artemisinins with the substrate pyridoxal inhibits PLP biosynthesis as demonstrated by kinetic measurements. Electrophysiological recordings from hippocampal slices and activity measurements of glutamic acid decarboxylase (GAD), a PLP-dependent enzyme synthesizing the neurotransmitter γ-aminobutyric acid (GABA), define how artemisinins interfere presynaptically with GABAergic signaling. Our data provide a comprehensive picture of artemisinin-induced effects on inhibitory signaling in the brain.

## INTRODUCTION

Pyridoxal 5’-phosphate (PLP) is the active form of vitamin B6. In humans, PLP biosynthesis is catalyzed by pyridoxal kinase (PDXK), a member of the ribokinase superfamily. PDXK utilizes inactive forms of vitamin B6 (pyridoxal (PL), pyridoxine and pyridoxamine) and ATP as substrates, producing PLP along with the byproduct ADP. The corresponding reaction proceeds *via* a random substrate addition reaction mechanism (Li et al., 2004) in which PLP biosynthesis takes place by transferring the γ-phosphate of ATP to the 5’-OH group of the B6 vitamers, in a process assisted by divalent metal ions such as Zn^2+^ and Mg^2+^ (Neary and Diven, 1970) **(Figure 1A)**. PLP serves as the essential active site component for more than160 distinct human enzymatic activities (di Salvo et al., 2012) catalyzing crucial cellular processes such as detoxification reactions and multiple metabolic processes including amino acid, carbohydrate and lipid metabolism. PLP-dependent enzymes also participate in neurotransmitter biosynthesis including the inhibitory neurotransmitters γ-aminobutyric acid (GABA) and glycine (di Salvo et al., 2012; Eliot and Kirsch, 2004; Percudani and Peracchi, 2003), which are synthesized by glutamic acid decarboxylase (GAD) and serine hydroxymethyl transferase (SHMT), respectively. Vitamin B6 deficiency has been implicated in multiple neurological, psychiatric and internal disorders possibly including even diabetes, cancer and autism (Merigliano et al., 2018), thus underpinning the importance of a finely tuned PLP biosynthesis.

**Figure 1.**
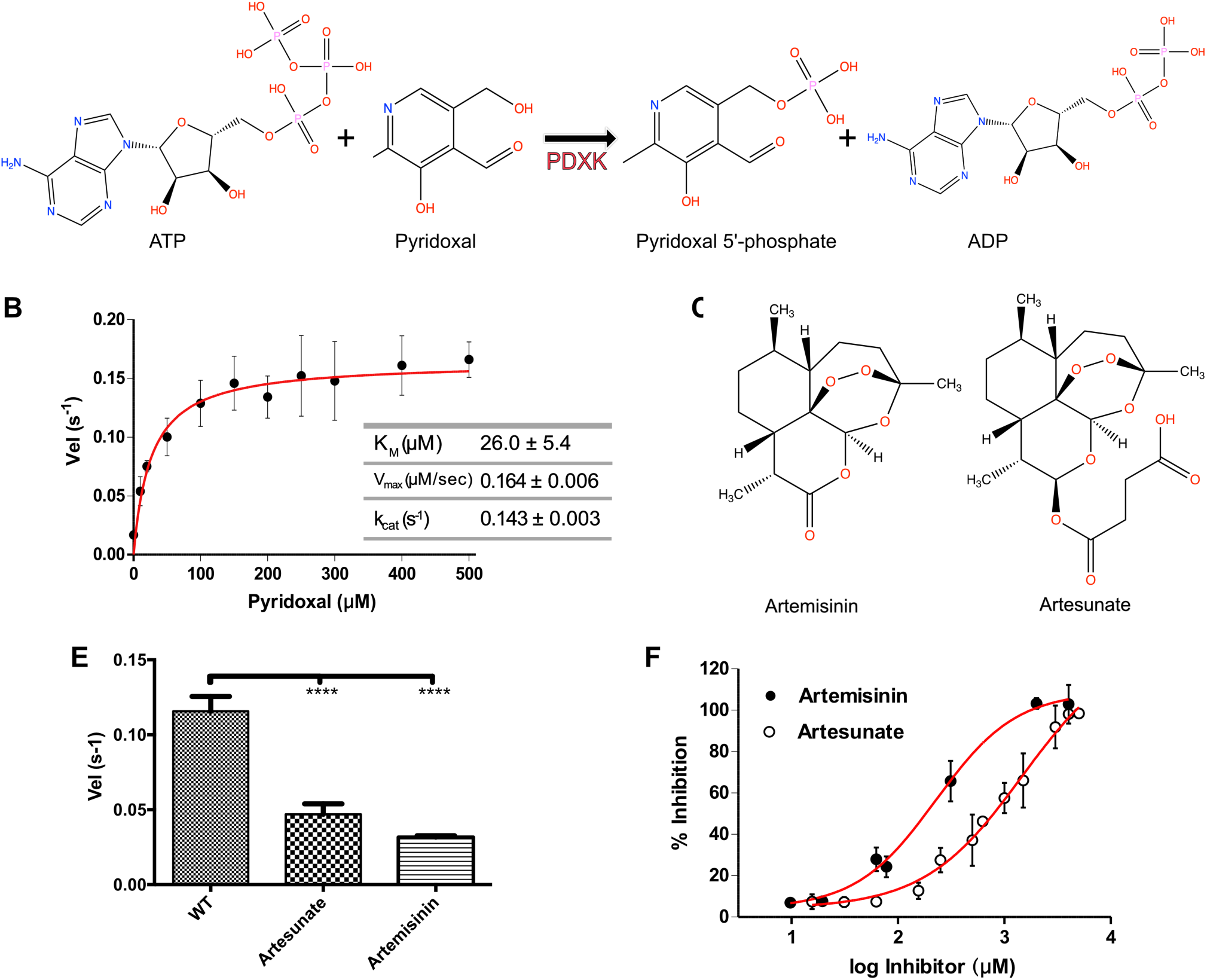
Biochemical Basis of PDXK Inhibition by Artemisinins. **(A)** Schematic representation of the reaction catalyzed by pyridoxal kinase (PDXK). **(B)** Michaelis-Menten curve derived for the enzymatic activity of recombinantly purified PDXK. **(C-D)** Chemical structures of artemisinin **(C)** and artesunate **(D)**. **(E)** Enzymatic activity of wild-type PDXK (WT-PDXK) in the absence and presence of artemisinin derivatives at a concentration of 1.5 mM (artesunate) and 156 μM (artemisinin), respectively. Data are presented as mean ± SEM. (p values are: *p<0.05; **p<0.01; ***p<0.001; ****p<0.0001) (Paired *t* test). **(F)** Inhibition curves of PDXK by artemisinin and artesunate used to derive the corresponding IC_50_ values.

Recently, PDXK was identified as one of the mammalian targets of the anti-malarial drug artemisinin (Li et al., 2017). Artemisinin-containing plant extracts have been used in traditional Chinese medicine for the treatment of malaria (Tu, 2016). Chemically, these small molecules are sesquiterpene lactones with an unusual endo-peroxide bridge. Artemisinin and its semi-synthetic derivatives artemether and artesunate (collectively referred to as artemisinins), in combination with quinones such as mefloquine and lumefantrine, nowadays represent the standard drug combinations used to treat malaria caused by *Plasmodium falciparum* (WHO, 2015). In addition to their anti-protozoan activities, these drugs have also been pharmacologically observed to regulate the activities of a variety of mammalian cellular processes some of which are deregulated in various types of cancer (Crespo-Ortiz and Wei, 2012; Gautam et al., 2009). Recently, it was discovered that artemisinins also modulate the differentiation of pancreatic Tα cells by inducing a trans-differentiation of glucagon-producing Tα cells into insulin-secreting Tβ cells, thus suggesting an anti-diabetic activity of artemisinins (Li et al., 2017). However, two subsequent studies contradicted this observation, thus questioning the potential clinical application of these compounds in the treatment of diabetes (Ackermann et al., 2018; van der Meulen et al., 2018).

Until recently, in the absence of a single protein crystal structure in complex with artemisinins (neither a plasmodial nor a mammalian protein), the detailed framework describing the target recognition by these small molecules remained enigmatic. The first molecular insights into artemisinin-recognition by a target protein were derived by us from crystal structures of the C-terminal domain of the moonlighting protein gephyrin (GephE) in complex with two artemisinin derivatives, artesunate and artemether (Kasaragod et al., 2019). Gephyrin is the principal scaffolding protein at inhibitory postsynaptic specializations and also catalyzes the final two steps of the evolutionarily conserved molybdenum cofactor (Moco) biosynthesis (Kasaragod and Schindelin, 2016, 2018; Kuper et al., 2004). Structures of the GephE-artemisinin complexes demonstrated that artemisinins specifically target the universal receptor binding pocket of this moonlighting protein, without altering its enzymatic activity, thus inhibiting critical interactions of gephyrin with GABA type A receptors (GABA_A_Rs) and glycine receptors (GlyRs). As an important functional consequence, artemisinins modulate inhibitory neurotransmission in a gephyrin-dependent manner. In addition to gephyrin, various proteins were identified as putative targets of artemisinins in pancreatic cells, including the central metabolic enzyme PDXK (Li et al., 2017), yet the molecular mechanisms underlying the modulation of these targets by artemisinins remained enigmatic.

Here, we determined the 2.4 Å resolution crystal structure of mouse pyridoxal kinase (mPDXK) in complex with artesunate, a succinate derivative of artemisinin. The artesunate binding site partially overlaps with the substrate (PL)/product (PLP) binding site, thus suggesting a drug-induced inhibitory effect. Enzymatic activity assays *in vitro* indeed revealed a significant inhibition of PLP production in the presence of artemisinins with K_i_ values in the high micromolar range. Electrophysiological recordings and measurements of GABA biosynthesis suggests that artemisinins exert their effect by down regulating the activity of PLP-dependent enzymes such as GAD. Taken together, our data define the molecular basis for the inhibition of PDXK by artemisinins and their consequences at the presynaptic terminals of inhibitory postsynapses and extend our current understanding of the artemisinin-induced modulation of inhibitory neurotransmission beyond gephyrin.

## RESULTS

### Artemisinins Inhibit PDXK

To derive the oligomeric state of recombinantly purified mPDXK, we first performed multi-angle light scattering coupled to size exclusion chromatography (SEC-MALS) experiments. The experiments showed that the protein is a dimer in solution **(Figure 1 – figure supplement 1)**, as has been reported for the human and also prokaryotic PDXK homologs (Kerry et al., 1986). Next, to check the activity of the recombinant enzyme, we measured its enzymatic activity by directly monitoring PLP production in a photometric assay. The characterization of the recombinantly purified protein showed a K_M_ of 26.0 ± 5.4 μM, a V_max_ of 0.1640 ± 0.006 μM/s and a k_cat_ of 0.1436 ± 0.003 s^-1^ for the substrate PL in the presence of 1 mM of ATP **(Figure 1B)**, which is in line with reported K_M_ values (3-50 μM) for PDXK (Elsinghorst et al., 2015; Hanna et al., 1997; Jones et al., 2012; Kwok and Churchich, 1979; McCormick et al., 1961; Safo et al., 2006; Ubbink et al., 1990).

To understand the effect of artemisinins on the enzyme, we performed the activity assays in the presence of two artemisinins, the parental compound artemisinin and artesunate **(Figure 1C-D, Source data 1)**. The determination of the turnover rates (Vel) displayed a highly significant inhibition in mPDXK activity in the presence of artemisinins with observed reductions to 0.032 ± 0.001 and 0.047 ± 0.007 s^-1^ for artemisinin and artesunate, respectively. Compared to the turnover rate of the enzyme in the absence of these drugs (0.116 ± 0.01 s^-1^) **(Figure 1E)** this corresponds to a ∼3-fold decrease. The enzymatic or turnover velocity, Vel, is defined here as the mean number of product molecules generated by a single enzyme per unit time. Statistical analyses revealed a significant reduction in enzymatic activity in the presence of the artemisinins and via a Dixon plot analysis K_i_-values of 120 ± 2.4 and 1250 ± 4.7 μM were derived for artemisinin and artesunate, respectively **(Figure 1 – figure supplement 2, Source data 1)**. To further characterize the inhibitory properties of artemisinins we also determined the IC_50_ for both compounds. While artesunate displayed an IC_50_-value of 1445 ± 1.4 μM, artemisinin was ∼6-fold more potent with an IC_50_ of 229 ± 1.3 μM **(Figure 1F)**.

### Structural Basis for the Inhibition of PDXK by Artemisinins

To gain insights into the mechanism of inhibition at the atomic level we determined three crystal structures of mPDXK. First, we derived the crystal structure of mPDXK in its apo-state and in complex with ATPγS, one of the substrates of the enzyme. These structures were solved by molecular replacement (MR) with the structure of human PDXK in the absence of any substrate as search model. The apo and the mPDXK-ATPγS structures were refined in space group C2 containing two dimers in the asymmetric unit to resolutions of 2.45 and 2.9 Å, respectively **(Table 1, Figure 2 – figure supplement 1A-B)**. The overall architecture of mouse apo-PDXK shares high structural similarity with its human ortholog (Li et al., 2002; Musayev et al., 2007) as reflected in root mean square (rms) deviations of 0.84 Å (PDB: 2YXT; human apo-PDXK) after superposition of all C_α_-atoms.

**Table 1.** Data Collection and Refinement Statistics.

|  | PDXK-apo | PDXK-ATP<sub>γ</sub>S | PDXK-ATP<sub>γ</sub>S-artesanate |
| --- | --- | --- | --- |
| **Data collection** |  |  |  |
| Space group | C2 |  |  |
| *a*, *b*, *c* (Å) | 279.13, 53.43, 109.37 | 278.60, 53.02, 109.85 | 279.38, 53.04, 110.15 |
| *α*, *β*, *γ* (°) | 90, 90.00, 90 | 90, 91.75, 90 | 90, 91.64, 90 |
| Resolution (Å) | 47.32 - 2.45 (2.53-2.45) | 47.16 - 2.9 (3.03-2.9) | 47.20- 2.4 (2.46-2.4) |
| <sup>a</sup>*R*<sub>sym</sub> | 0.098 (0.75) | 0.10 (0.739) | 0.084 (1.069) |
| <sup>b</sup>*R*<sub>pim</sub> | 0.093 (0.70) | 0.068 (0.488) | 0.053 (0.704) |
| *CC*<sub>1/2</sub> | 0.993 (0.572) | 0.995 (0.600) | 0.996 (0.541) |
| <sup>c</sup><\|*I*/*σ*\|> | 9.1 (1.6) | 9.3 (1.6) | 8.1 (1.1) |
| Completeness (%) | 95.6 (99.7) | 99.7 (99.7) | 99.7 (99.6) |
| Redundancy | 3.2 (3.5) | 3.4 (3.2) | 3.4 (3.1) |
| Reflections used in refinement | 57174 (5965) | 36057 (3532) | 63558 (6235) |
| **Refinement** |  |  |  |
| <sup>d</sup>*R*-work | 0.2162 (0.3042) | 0.2341 (0.3378) | 0.2088 (0.3271) |
| <sup>e</sup>*R*-free | 0.2596 (0.3237) | 0.2548 (0.3366) | 0.2535 (0.3851) |
| Number of non-hydrogen atoms | 9715 | 9862 | 9890 |
| Macromolecules | 9411 | 9606 | 9445 |
| Ligands | 116 | 232 | 304 |
| Solvent | 188 | 24 | 141 |
| RMS(bonds) | 0.002 | 0.005 | 0.002 |
| RMS(angles) | 0.53 | 0.73 | 0.54 |
| <sup>f</sup>Ramachandran favored (%) | 96.83 | 96.99 | 96.76 |
| Ramachandran allowed (%) | 3.17 | 3.01 | 3.24 |
| Ramachandran outliers (%) | 0.00 | 0.00 | 0.00 |
| Average B-factor (Å<sup>2</sup>) | 57.79 | 95.87 | 69.33 |
| Protein | 57.62 | 95.89 | 68.94 |
| Ligands | 74.74 | 95.54 | 84.27 |
| Solvent | 55.93 | 91.85 | 63.08 |

<sup>a</sup>*R*<sub>sym</sub> = $\sum_{hkl} \sum_i |I_i - \langle I \rangle| / \sum_{hkl} \sum_i I_i$ where *I<sub>i</sub>* is the *i*<sup>th</sup> measurement and *⟨I⟩* is the weighted mean of all measurements of *I*.
<sup>b</sup>*R*<sub>pim</sub> = $\sum_{hkl} 1 / (N-1)^{1/2} \sum_i |I_i(hkl) - \overline{I(hkl)}| / \sum_{hkl} \sum_i I_i(hkl)$ , where N is the redundancy of the data and
$\overline{I(hkl)}$ the average intensity.
$^c\langle I/\sigma \rangle$ indicates the average of the intensity divided by its standard deviation.
$^dR_{\text{work}} = \sum_{hkl} ||F_o| - |F_c|| / \sum_{hkl} |F_o|$ where $F_o$ and $F_c$ are the observed and calculated structure factor amplitudes.
$^eR_{\text{free}}$ same as R for 5% of the data randomly omitted from the refinement. The number of reflections includes the $R_{\text{free}}$ subset.
$^f$ Ramachandran statistics were calculated with MolProbity.
Although the $\beta$ angle of the apo structure is 90.00 after post-refinement, the crystals display monoclinic and not orthorhombic symmetry.
Numbers in parentheses refer to the respective highest resolution data shell in each dataset.

Closer inspection of the nucleotide binding pocket revealed that ATPγS-binding is directly mediated by Val226, which forms a hydrogen bond with the adenine of the nucleotide through its main chain carbonyl oxygen and residues Thr186 and Thr233 as well as Asp118, Asn150, which coordinate the ATP analog through interactions with the α and β-phosphate of its tri-phosphate moiety, respectively **(Figure 2 – figure supplement 1C)**. There were no significant conformational changes in the binary complex compared to the mouse apo structure as reflected in an rms deviation of 0.45 Å for all C_α_-atoms with minimal structural rearrangements in ATP binding pocket **(Figure 2 – figure supplement 1D)**. Comparison of PDXK sequences derived from organisms representing different evolutionary levels revealed that all residues, which are crucial for the binding of the nucleotide, are strictly conserved **(Figure 2 – figure supplement 2)**.

To gain insights into the mechanism of artemisinin inhibition we determined the crystal structure of mPDXK in complex with ATPγS and artesunate **(Figure 2, Table 1)**. This structure was obtained by soaking artesunate into pre-existing binary mPDXK-ATPγS crystals. After molecular replacement with the apo structure, in addition to the clear density for ATPγS **(Figure 2B)**, strong difference density in close proximity to the substrate-binding pocket was also observed **(Figure 2D)**, which allowed us to unambiguously model the bound artesunate. Surprisingly, this density was observed in only one of the four molecules present in the asymmetric unit. The absence of artesunate in the other monomers may be due to the involvement of these protomers in crystal contacts, thus preventing artesunate-binding when soaking the compounds into pre-existing crystals.

**Figure 2.**
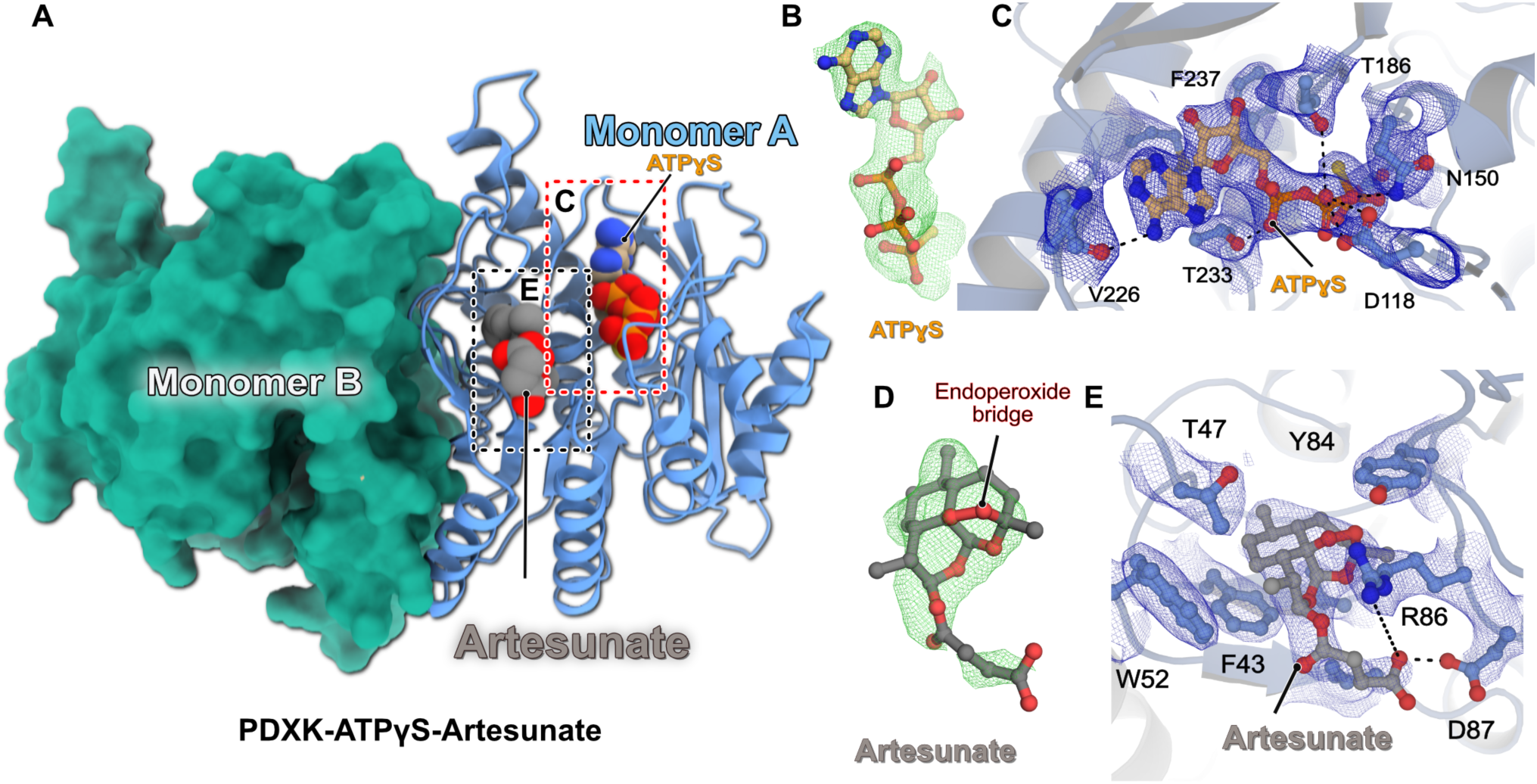
Structure of the Ternary PDXK-ATPγS-Artesunate Complex. **(A)** Overall architecture of the ternary complex. One monomer is shown in cartoon representation with the bound ligands in CPK representation, while the second monomer is shown as a surface in green. **(B)** F_o_-F_c_ omit electron density for the bound ATPγS contoured at an rms deviation of three. **(C)** Enlarged view of the ATPγS binding pocket. The bound ligand and residues, which are crucial for ligand binding, are shown in ball-and-stick representation. The SIGMAA-weighted 2F_o_-F_c_ electron density for the bound ligand and surrounding residues is contoured at an rmsd of one. Critical protein-ligand interactions are highlighted. **(D)** F_o_-F_c_ omit electron density for the bound artesunate contoured at an rms deviation of three. **(E)** Enlarged view of the artesunate-binding pocket. The bound ligand and residues, which are crucial for ligand binding, are shown in ball-and-stick representation. SIGMAA-weighted 2F_o_-F_c_ electron density for artesunate and interacting residues contoured at an rmsd of one. Critical protein-ligand interactions are highlighted.

The fact that all three structures reported here belong to the same space group with similar unit cell parameters and essentially identical crystal packing allowed for a meaningful comparative analysis. The overall architecture of the mPDXK-ATPγS-artesunate structure is identical to the apo and binary mPDXK-ATPγS structures; a superposition of the C_α_ atoms of these two complexes revealed rms deviations of 0.51 and 0.30 Å for the apo and ATPγS-bound structures, respectively. Binding of the substrate analog ATPγS was mediated by the same residues described for the binary mPDXK-ATPγS complex **(Figure 2C and Figure 2 – figure supplement 3)**. A closer inspection of the artesunate-binding pocket revealed that drug-binding is mainly mediated by Val41, Thr47 and also Trp52, which generate a hydrophobic pocket that binds artesunate binds with a buried surface area of 364 Å^2^ compared to a total surface area of the drug of 538 Å^2^. In particular, artesunate is sandwiched in between two aromatic residues, Phe43 and Tyr84, which stabilize artesunate through van der Waals interactions. In addition to the hydrophobic interactions, the carboxylate moiety of artesunate comes into proximity of the guanidinium group in the side chain of Arg86, which potentially stabilizes the interactions through electrostatic contacts. Finally, Asp87 favors artesunate-binding through a hydrogen bond (2.5 Å) between its side chain and the carboxylate of the artesunate assuming one of these carboxylates is protonated **(Figure 2E)**.

An analysis of the mPDXK-ATPγS-artesunate structure showed that the ATP binding pocket is in relatively close proximity from the bound artesunate at a distance of ∼21 Å, as measured between the C_α_ atoms of Phe237 in the ATP binding pocket and Phe43 in the artesunate binding pocket. A comparison of the ternary mPDXK-ATPγS-artesunate and binary mPDXK-ATPγS structures revealed that binding of artesunate neither induced any significant rearrangements in the conformation of the nucleotide nor in the residues mediating its binding, in line with the independent binding of the two ligands **(Figure 2 – figure supplement 3 and 4)**. Interestingly, the binding pocket of artesunate, like that of ATP as mentioned earlier, is highly conserved **(Figure 2 – figure supplement 5)**. To get additional information regarding the mode of inhibition, we also compared our ternary structure with the PDXK-PLP structures. Strikingly, when we superimposed the already reported ternary *Hs*PDXK-ATP-PLP (PDB: 3KEU) structure with our ternary complex, a critical partial overlap between the tri-cyclic ring system of artesunate and the pyridine ring of the product PLP was uncovered **(Figure 3A-B)** which would result in severe van der Waals repulsions for the C2 and C3 atoms of PLP and the C2 and C3 atoms of artesunate, if bound simultaneously. This clearly constitutes a major reason for the inhibition of the PDXK. Moreover, an analysis of the surface properties of the ternary structure revealed a tunnel, which is leading from the protein surface to the distal end of ATP-binding pocket spanning a length of ∼38 Å, which is blocked near its entrance by artesunate. A blockade of this tunnel, in turn, may prevent an efficient turnover of the enzyme. Taken together, these data illustrate the structural basis for the inhibition of mPDXK by artemisinins **(Figure 3C-E)**.

**Figure 3.**
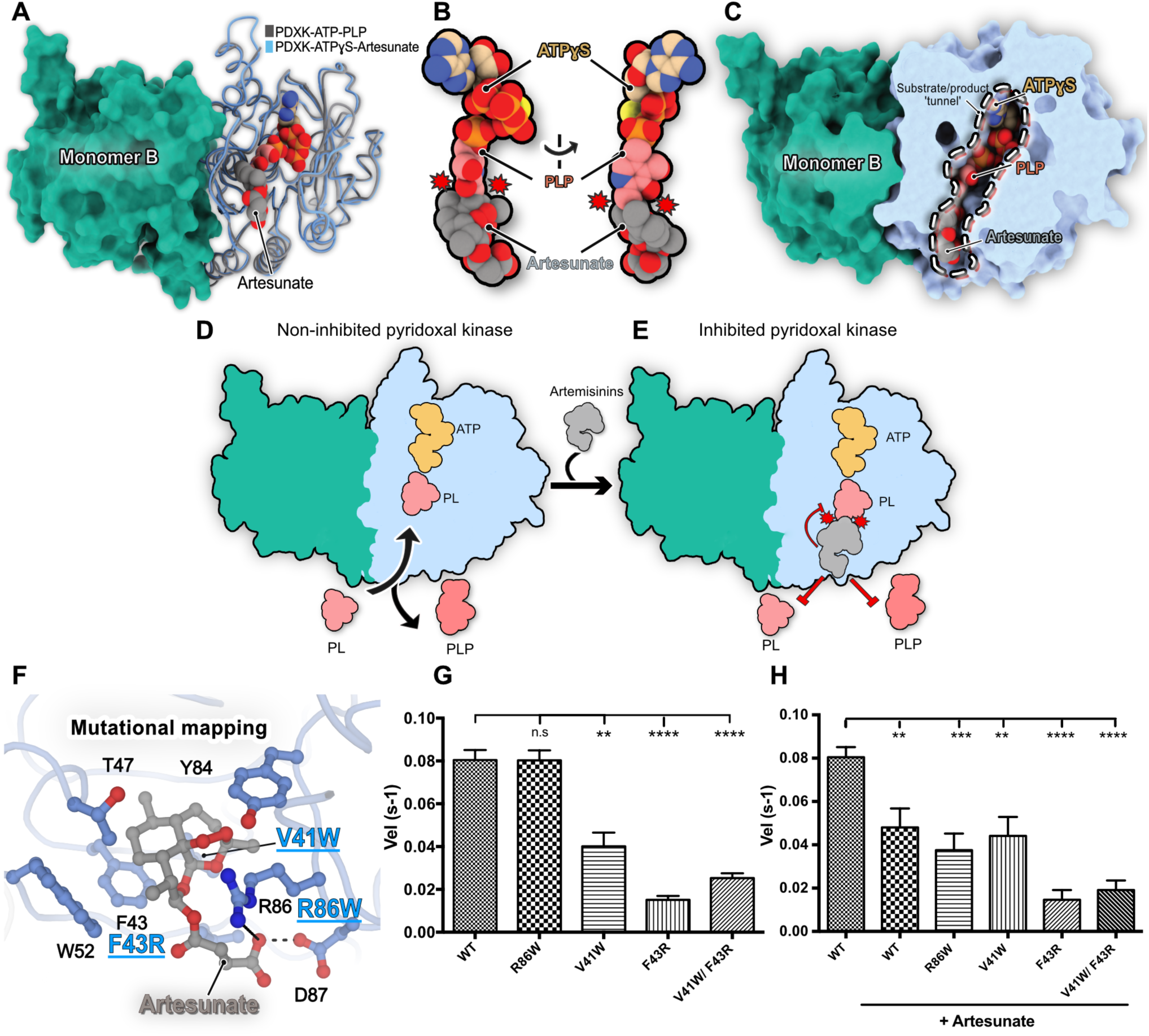
Structural basis for PDXK Inhibition by Artemisinins. **(A)** Superposition of the crystal structures of the ternary murine PDXK-ATPγS-artesunate (this study) and the human PDXK-ATP-PLP (PDB entry 3KEU) complexes. Carbon-atoms of artesunate are shown in gray, those of PLP in pink and those of ATPγS in beige, the other atoms are colored in red (oxygen), blue (nitrogen), orange (phosphorous) and yellow (sulfur). Only ATPγS of the artesunate complex is shown to reduce visual complexity. **(B)** Enlarged view of the ligand binding pockets of ATP, artesunate and PLP displaying the partial overlap (indicated by red circles with spikes) between PLP and artesunate. **(C)** Cut-away view of the superimposed PDXK-ATPγS-artesunate and PDXK-ATPγS-PLP structures displaying how artesunate binding blocks the substrate tunnel which may also impair enzyme turnover. **(D-E)** Schematic representation of structural basis for the inhibition of PDXK activity by artemisinins. **(F)** Mutational mapping of the artesunate-binding pocket with the investigated mutants highlighted in blue. **(G-H)** Comparison of the turnover rates of PDXK variants and the WT in the absence **(G)** and presence **(H)** of artesunate. (*p<0.05; **p<0.01; ***p<0.001; ****p<0.0001) (Paired *t*-test).

### Mapping of the Artemisinin Binding Pocket

To validate the observations derived from the crystal structures, we performed site directed mutagenesis experiments of residues located in the artesunate binding pocket **(Figure 3F)** and tested these mutants for PLP production in the presence and absence of artesunate **(Figure 3G-H, Source data 2)**. First, we analyzed the mutants through SEC-MALS, which revealed that all variants retained their dimeric state in solution as observed for the wild-type (WT) protein **(Figure 3 – figure supplement 1)**. Amongst the residues being investigated, we mutated Val41 and Phe43, which are involved in mediating the binding of artesunate through hydrophobic interactions to either introduce steric interference or alter the polar properties of the binding pocket, respectively. The V41W and F43R mutants significantly lowered the turnover rates (0.04 ± 0.006 s^-1^ for V41W and 0.016 ± 0.001 s^-1^ for F43R), even in the absence of artesunate in comparison to the WT (0.080 ± 0.004 s^-1^). This can be easily explained by the fact that these residues also mediate binding of PL and thus play a role in the regular enzymatic turnover of the protein. In contrast, mutation of Arg86, the residue involved in the long range electrostatic interaction with the carboxylate of artesunate to the bulky aromatic side chain of Trp did not alter the activity of mPDXK in the absence of artesunate, as demonstrated by its turnover rate of 0.08 ± 0.005 s^-1^, which is virtually identical to that of the WT **(Figure 3G)**.

Next, we further analyzed the catalytic activity of the mutants in the presence of artesunate **(Figure 3H and Figure 3 – figure supplement 2)**. The V41W and F43R variants did not result in significant changes in the turnover rates of the enzyme in the presence of artesunate (0.044 ± 0.009 for V41W and 0.014 ± 0.005 for F43R s^-1^ compared to 0.040 ± 0.006 and 0.016 ± 0.001 s^-1^, respectively, in its absence), which is in line with artesunate binding being abolished in both variants **(Figure 3 – figure supplement 2C and D)**. As expected, a similar trend was observed in case of the V41W/F43R double mutant with turnover rates of 0.024 ± 0.002 s^-1^ in the presence of artesunate compared to 0.019 ± 0.004 s^-1^ in its absence **(Figure 3 – figure supplement 2E)**. An identical behavior was observed in the case of GephE where mutation of a crucial aromatic residue (Phe330) to Ala completely abolished artemisinin binding (Kasaragod et al., 2019). In contrast, when we compared the activity of the R86W variant in the absence (0.080 ± 0.005 s^-1^) and presence (0.038 ± 0.008 s^-1^) of the drug, a significant reduction in enzymatic activity was observed **(Figure 3 – figure supplement 2B)**. Thus, although R86 is involved in an electrostatic interaction as revealed by the crystal structure, the mutational analysis demonstrated that the inhibition potency of artesunate is retained even in the absence of this interaction. This observation is in contrast to the GephE structure where the replacement of Arg (Arg653 in gephyrin) with the bulkier aromatic Trp, prevented artesunate binding (Kasaragod et al., 2019). Thus, our structures help to define the molecular signatures of artemisinin-binding pockets, which may aid in the future identification of target sites, especially by *in silico* approaches.

### Artemisinins inhibit GABA biosynthesis and downregulate GABAergic neurotransmission

To understand if the functional consequences of our biochemical and structural analyses correlate with a physiological scenario, we performed whole-cell voltage-clamp recordings (**Figure 4, Source data 3)** from CA1 pyramidal cells in hippocampal slices and determined the properties of GABAergic miniature inhibitory postsynaptic currents (mIPSCs) in the absence and presence of artemisinins (10 and 30 μM, **Figure 4A-B**), measured within 10 minutes of drug application. In line with our earlier findings (Kasaragod et al., 2019), artemisinin down-regulated mIPSC amplitudes already at 10 μM (from 56.7 ± 1.9 pA to 38.8 ± 2.6 pA, n = 7 from 4 mice, p = 0.003, paired t-test; **Figure 4A and C**), which we attribute to the artemisinin-induced disruption of postsynaptic GABA_A_R-gephyrin complexes. In addition, artemisinin altered mIPSC kinetics with slower rise and decay times **(Figure 4E-F)**. While a significant reduction in amplitudes was retained at a higher concentration (30 μM), we also observed a concomitant decrease in mIPSC frequency from 5.0 ± 0.61 Hz to 3.9 ± 0.48 Hz, (n = 7 from 5 mice, p = 0.003, **Figure 4B and D**) in the presence of artemisinin. This is particularly noteworthy in the context of this study as it reflects changes in the presynaptic terminals, e.g. it would be in line with a reduced synthesis of the neurotransmitter GABA (Engel et al., 2001).

**Figure 4.**
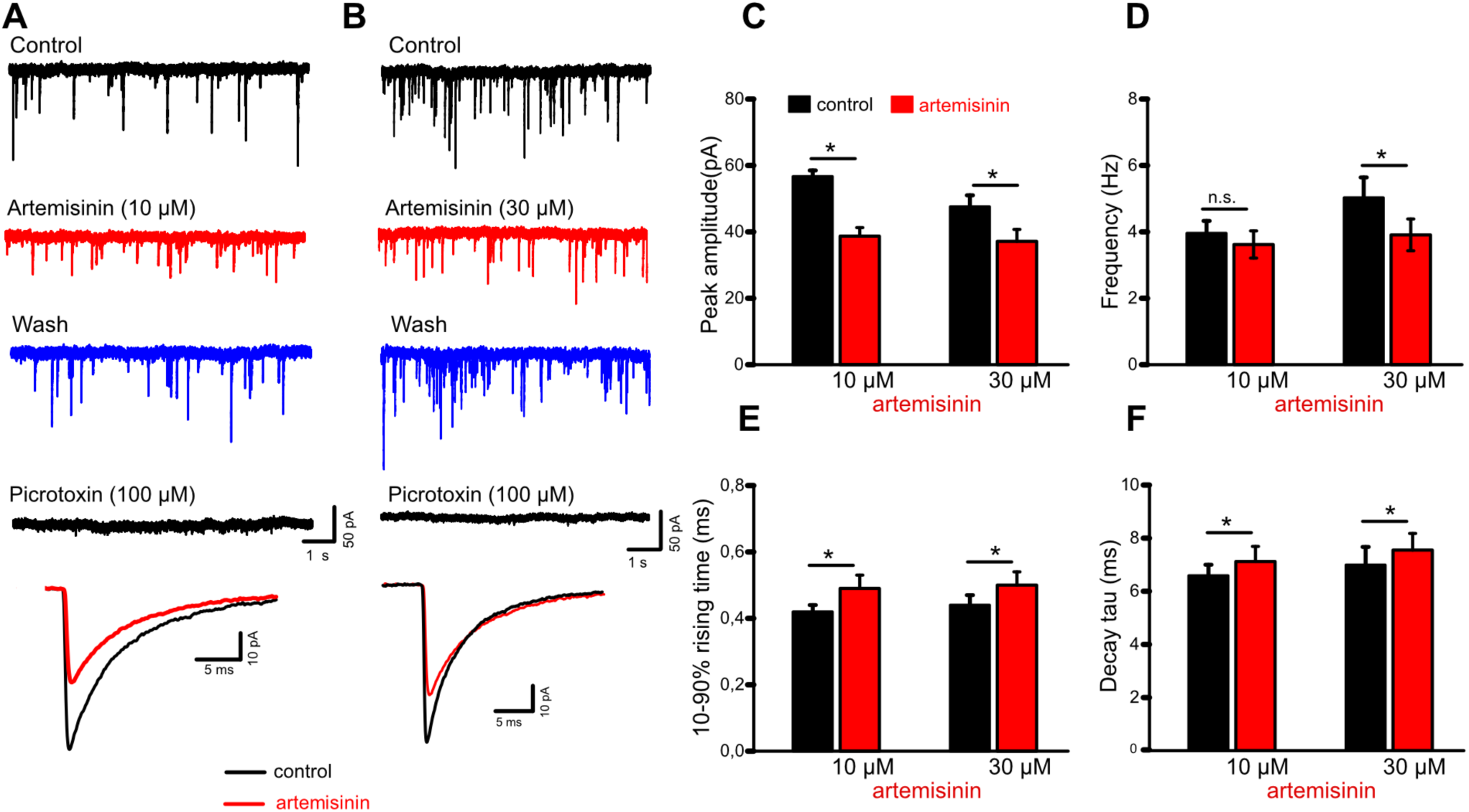
Impact of Artemisinins on Electrophysiological Recordings of Hippocampal Slices: **(A-B)** Representative voltage-clamp recordings of GABA_A_ receptor-mediated mIPSCs from mouse CA1 pyramidal cells in hippocampal slices were collected before (control), during (10 mins) and after (wash) artemisinin (10 µM, **A** and 30 µM, **B**) application. Picrotoxin was applied to verify the GABAergic origin of these events. Superimposed traces below are averaged events before and during artemisinin treatment. (**C-D**) Quantifications of how artemisinin treatment affects mIPSC amplitudes (**C**), frequencies (**D**) and kinetics (**E-F**) (10 µM: n = 7; 30 µM: n = 7). * p < 0.05, paired *t-*test.

To evaluate if the decreased mIPSC frequencies were due to changes in the activity of GAD, the GABA synthesizing enzyme, we first quantified the expression of this enzyme in hippocampal neurons (DIV14 **Source data 4**). This analysis revealed no altered protein expression levels **(Figure 5A and B)**. Next, we analyzed the expression levels of PDXK **(Figure 5A and C)**, which synthesizes the obligatory cofactor PLP for the activity of GAD and observed that its expression level was also unaffected by artemisinin treatment. As a qualitative measure, we also stained hippocampal neurons for PDXK, GAD and the postsynaptic marker gephyrin, which did not reveal any noticeable differences in the expression of these marker proteins **(Figure 5 – figure supplement 1 and 2)**. Finally, to check if the frequency changes observed in the electrophysiology measurements were due to changes in the activity of GAD, we measured the amount of GABA being synthesized in primary hippocampal neurons (DIV14). GABA levels were quantified by the classical ninhydrin reaction **(Figure 5 – figure supplement 3)** by measuring the fluorescence emission of the resulting adduct at 450 nm and calibrating it with a GABA standard. Remarkably, this analysis revealed a significant reduction in the amount of GABA production (5.4 ± 0.8, 3.2 ± 0.9 and 3.6 ± 0.98 GABA/mg protein/h) in hippocampal neurons treated with artemisinin at concentrations of 3, 10 and 30 μM, respectively (p=0.0054, 0.0007 and 0.0002 against DMSO measurements and p=0.018, 0.0087 and 0.0047 against hippocampal measurements for 3, 10 and 30 μM artemisinin concentrations) **(Figure 5D)**. In comparison, untreated samples resulted in levels of 9.8 ± 1.5 µg GABA/mg protein/h. The observed perturbation on the presynaptic side is therefore due to the direct effect of artemisinins on the biosynthesis of PLP, which results in a reduced production of this cofactor, which is required by the GAD enzyme to produce the neurotransmitter GABA **(Figure 5E)**.

**Figure 5.**
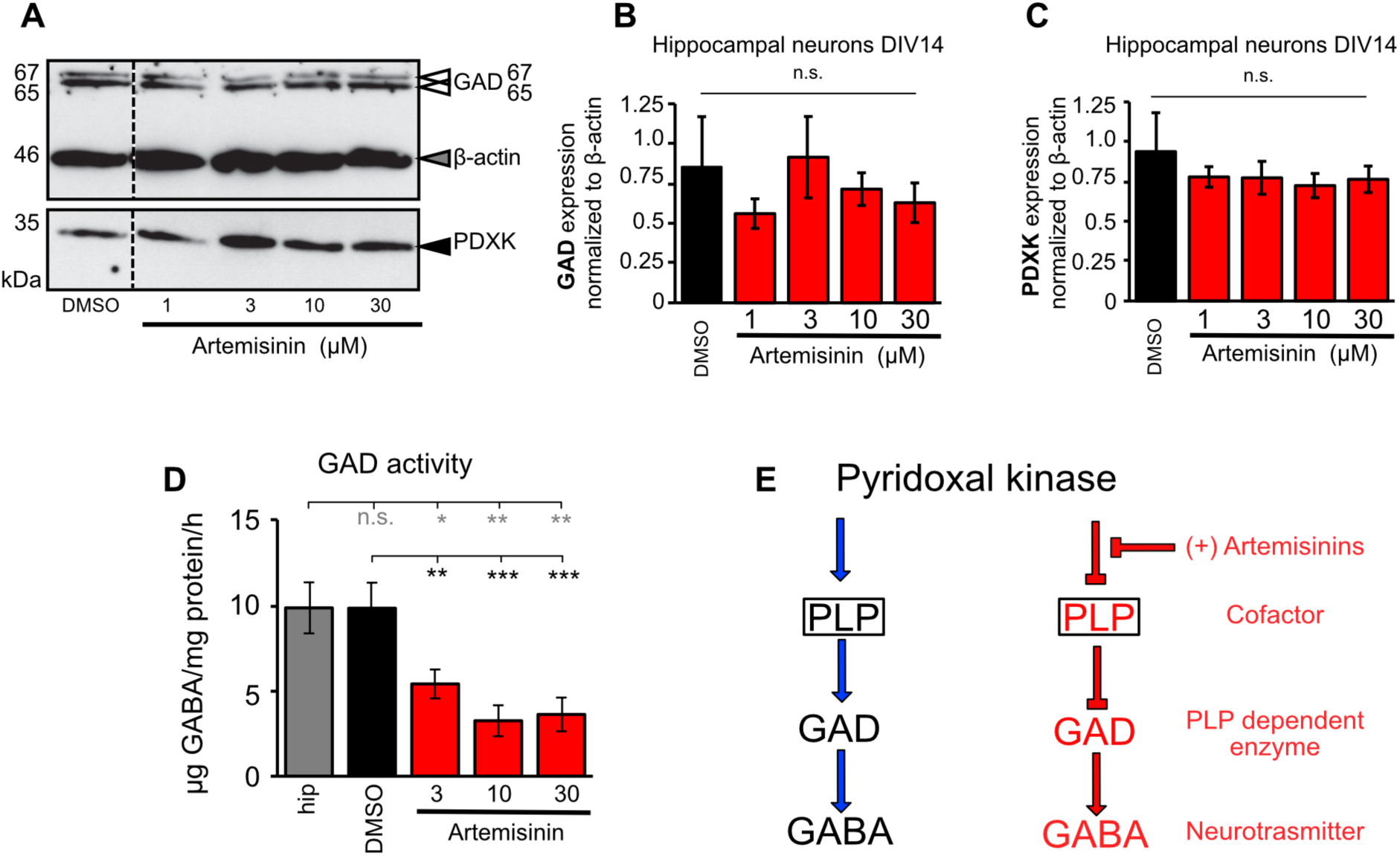
Artemisinins impact GABA biosynthesis by down regulating GAD activity: **(A)** Representative image of a Western blot stained for GAD (GAD isoforms at 65 and67 kDa are indicated by white arrowheads), PDXK (35 kDa, black arrowhead), and ß-actin (46 kDa, gray arrowhead). DMSO treated cells served as internal control. **(B)** Quantitative analysis of GAD from lysates of hippocampal neurons DIV14 incubated with increasing concentrations of artemisinin (1, 3, 10, 30 µM; DMSO served as control) for 2 h. Expression of GAD was normalized to β-actin expression. **(C)** PDXK expression analysis after incubation with artemisinins (1, 3, 10, 30 µM). PDXK expression was also normalized to ß-actin. **(D)** Measurements of GAD activity in hippocampal samples following treatment with different concentrations of artemisinin (3, 10, 30 µM). Tissue without treatment (gray bar) and treated with DMSO (black bar) served as positive controls. Number of measurements n = 8-9 from three independent biological replicates. GAD activity decreased significantly with increasing concentrations of artemisinin. The data were analyzed with a paired *t*-test. p=0.0054, 0.0007 and 0.0002 against DMSO measurements and p=0.018, 0.0087 and 0.0047 against hippocampal measurements for artemisinin concentrations of 3μM, 10μM and 30 μM. **(E)** Schematic representation of the steps leading to GABA biosynthesis at presynaptic terminals. The left panel shows how GAD synthesizes GABA by utilizing PLP as a cofactor which is produced by PDXK, while the right panel shows how artemisinins inhibit the initial step in the biosynthesis by inhibiting PDXK, which, in turn, indirectly impacts downstream biosynthetic processes and eventually downregulates the amount of neurotransmitter being synthesized.

## DISCUSSION

Despite their widespread clinical application as anti-malarial drugs, and despite their known effects on various cellular pathways in mammals, the molecular mechanisms of how artemisinins affect cellular pathways are still only poorly understood. Artemisinins can efficiently cross the blood-brain barrier (Davis et al., 2003) and, strikingly, administrations of high levels of artemisinins are accompanied by severe neurotoxic side effects (Brewer et al., 1994; Schmuck et al., 2002; Wesche et al., 1994). Recently, we were able to derive the first protein-artemisinin structure by X-ray crystallography at 1.5 Å resolution, namely that of the scaffolding protein gephyrin in complex with artesunate and artemether (Kasaragod et al., 2019). Here, we successfully validated and elucidated the mechanism underlying yet another mammalian artemisinin target, the critically important metabolic enzyme PDXK.

Our structural studies demonstrate a competition between the substrate pyridoxal and artemisinins, in line with the observed inhibition of the enzyme derived from kinetic data. As artesunate targets the same binding pocket identified previously for the interaction of (*R*)-roscovitine with PDXK (Bach et al., 2005; Tang et al., 2005) and for the neurotoxins ginkgotoxin and theophylline (Gandhi et al., 2012), our structure suggests that the neurotoxicity induced by artemisinins could be due, at least in part, to their binding to PDXK and the resulting inhibition of its activity.

The presynaptic effect of artemisinin in our electrophysiological recordings correlates nicely with the down-regulation of PDXK activity and can be extended towards glycine, the other major inhibitory neurotransmitter. This neurotransmitter is synthesized by serine hydroxymethyl transferase (SHMT), again in a strictly PLP-dependent fashion. Thus, we predict a similar electrophysiological behavior with decreased frequencies at glycinergic synapses as observed for GAD and GABA levels. We have already demonstrated a decrease in glycinergic currents following artemisinin treatment (Kasaragod et al., 2019). Thus, the data presented here extend our current understanding of how artemisinins act at inhibitory synapses in the CNS. The present study shows that artemisinins not only act at the postsynaptic side, but also affect the functionality of the presynaptic terminals via their interaction with PDXK which ultimately leads to a decrease in neurotransmitter biosynthesis **(Figure 6)**. Although the data presented here and our earlier study (Kasaragod et al., 2019) define mechanisms underlying the downregulation of inhibitory neurotransmission by artemisinins, other neurotransmitters such as dopamine, histamine and serotonin are also synthesized in a PLP-dependent manner, thus future studies will be required to comprehensively dissect the molecular details underlying the artemisinin-induced regulation of neurotransmitter levels and the resulting physiological consequences.

**Figure 6.**
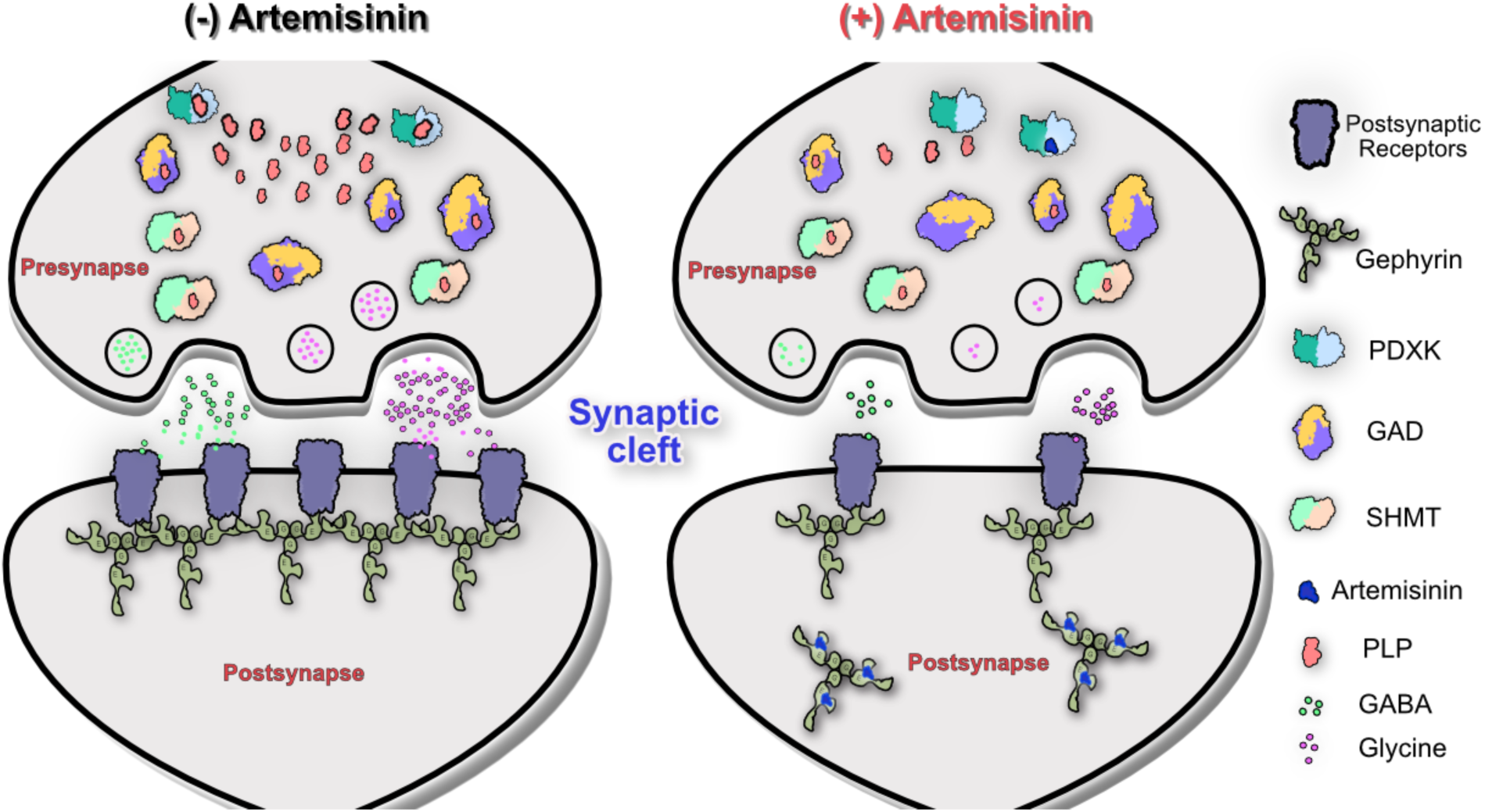
Schematic representation of inhibitory synapses in the absence and presence of artemisinins. This scheme shows that in the absence of artemisinins (left panel) gephyrin clusters the receptors required for inhibitory neurotransmitters at postsynaptic sides, while PDXK contributes to the biosynthesis of neurotransmitters at presynaptic terminals by producing the PLP cofactor for GAD and SHMT enzymes. In contrast, in the presence of artemisinins (right panel), gephyrin-mediated clustering of receptors at postsynaptic sites is impaired while neurotransmitter biosynthesis at presynaptic terminals is inhibited.

In addition, a comparison of the artemisinin binding pockets in mPDXK and GephE revealed common denominators of drug recognition. Notably, in both cases, artemisinins engage in crucial van der Waals interactions with aromatic residues. In addition, in GephE as well as PDXK, the side chain of an Arg contributes to artesunate binding, which stabilizes the drug through an electrostatic interaction with its succinate moiety, thus revealing common signatures of artemisinin binding pockets. Our results thus not only broaden the understanding of target recognition by artemisinins at the structural level, but also provide important new insights into how these interactions impair inhibitory synaptic transmission in the brain and on how this might account for the neurological side effects of these drugs. Future studies, along with molecular signatures revealed in our structures, will be required to investigate if artemisinins indeed directly bind to and also modulate the activities of other mammalian targets such as protein disulfide isomerase and fatty acid synthase which were identified earlier (Li et al., 2017).

## ACKNOWLEDGEMENTS

We want to thank Dr. Antje Gohla for providing the cDNA of mPDXK as well as Dana Wegmann and Christine Schmitt for their excellent technical assistance. We also thank the beamline scientists at beamline ID23-A1, ESRF, Grenoble, for technical assistance during data collection. We are grateful to Dr. Kunimichi Suzuki for the critical reading of the manuscript. This work was supported by the Deutsche Forschungsgemeinschaft (SCHI425/8-2) and the Rudolf Virchow Center for Experimental Biomedicine to HS. APM was supported by the Graduate School for Life Sciences at the University of Würzburg. VBK is supported at the MRC LMB by an EMBO Long Term Fellowship (ALTF137-2019).

## AUTHOR CONTRIBUTIONS

Conceptualization, V.B.K. and H.S.; Methodology, V.B.K., A.P.M., NS., F.Z., and N.B.; Investigation, V.B.K., A.P.M., N.S., F.Z., and N.B.; Writing – Original Draft, V.B.K.; Writing – Review & Editing, V.B.K., C.V., C.A. and H.S.; Funding Acquisition, H.S.; Resources, C.A. C.V, and H.S.; Supervision, V.B.K., C.A., C.V., and H.S.

## MATERIALS AND METHODS

### Experimental model and subject details

For cloning purposes *E. coli* DH5α was used and the cells were grown on LB-agar plates and in LB liquid medium at 37°C. For recombinant protein expression *E. coli* SoluBL21™ cells were used. The cells were grown at 37°C initially and were further incubated at 30°C for 16-18 hours after induction. Transverse hippocampal slices (350 µm thick) were prepared from sevoflurane-anesthetized adult C57Bl/6J mice (2 – 4 months old) of either sex purchased from Charles River (Sulzfeld, Germany). Adult animals (12 weeks old male or female mice) were taken from the mouse strain CD1 (Strain code: 022, Charles River, Sulzfeld, Germany) to isolate the hippocampi. Animals were housed under standard conditions and all procedures were conducted according to the guidelines and with approval of the local government. Preparation of brain slices containing the hippocampal formation was performed as previously described (Zheng et al., 2016)

### Method detail

#### Cloning, Recombinant Protein expression and Purification

The cDNA encoding mPDXK was subcloned into the pETM14 expression vector harboring a 3C-precision protease cleavage and BamH1 sites by sequence independent ligation cloning (SLIC) (Li and Elledge, 2007). The proteins (WT and all mutants) were expressed in the *E. coli* SoluBL21™ strain. Cells were grown at 37 °C and expression was induced with 0.5 mM IPTG at an optical density (OD_600_) of 0.6-0.8 and cultures were subsequently incubated at 30 °C for 16-18 hours. Following centrifugation at 8000 x *g* for 15 min the harvested cells were re-suspended in lysis buffer containing 50 mM Tris pH 8, 300 mM NaCl and 5 mM β-mercaptoethanol (β-ME) and lysis was performed by using a microfluidizer. For purification, a two-step protocol was employed consisting of an initial Ni-affinity chromatography with Ni-IDA beads, which was followed by cleavage of the N-terminal His_6_-tag by incubating with 3C precision protease overnight at 4°C. Finally, size exclusion chromatography on a Superdex 200 26/60 (GE Healthcare) column was performed in SEC buffer (20 mM Tris pH 8, 150 mM NaCl and 5 mM β-ME) to purify the protein to apparent homogeneity.

#### Size Exclusion Chromatography Coupled to Multi-Angle Light Scattering (SEC-MALS)

SEC-MALS experiments of 100 μM WT and all mutants were carried out by using a Superdex 200 10/300 column (GE Healthcare) in SEC buffer. The experiments were performed at a constant flow rate of 0.5 ml/min at room temperature. The differential refractive index (dRI) and the light scattering (LS) were monitored with a Dawn Helios detector from Wyatt Technologies and molecular masses were derived from the dRI and LS measurements.

#### Crystallization

Crystallization of mPDXK was performed in the apo form and in complex with ATPγS at a protein concentration of 12 mg/ml corresponding to a molar concentration of 0.3 mM. The protein was mixed with 2 mM ATPγS and 5 mM MgCl_2_ and the complex was incubated on ice for 30 min prior to crystallization. Crystallization was performed with the sitting drop vapor diffusion method by mixing equal volumes of protein and mother liquor at 20 °C. The mPDXK-ATPγS-artesunate structure was determined by soaking mPDXK-ATPγS crystals with different concentrations of artesunate (2-10 mM) for 30-600 sec. Crystals were transferred into mother liquor (0.18-0.24 M sodium thiocyanate and 18-26% PEG3350) supplemented with different concentrations of artesunate and 25% glycerol as cryo-protectant before flash cooling in liquid nitrogen.

#### Data Collection, Structure Determination and Refinement

Data collection for all crystals was performed at the ESRF, Grenoble, on beamline ID23A-1 at a wavelength of 0.9724 Å at 100 K. Datasets were indexed and integrated with XDS (Kabsch, 2010) and subsequently scaled and merged with AIMLESS (Evans and Murshudov, 2013) from the CCP4 suite (Winn et al., 2011). The apo-structure and the binary mPDXK-ATPγS complex were determined by molecular replacement with PhaserMR (McCoy et al., 2007) using the human PDXK structure (PDB: 2YXT) as search model and the ternary mPDXK-ATPγS-artesunate complex was solved with the apo mPDXK structure as search model. The protein crystallized in space group C2 with four molecules in the asymmetric unit. Refinement was performed in PHENIX (Adams et al., 2010) with repeated manual model building in COOT (Emsley and Cowtan, 2004). Coordinates and restraints for artesunate were obtained from our gephyrin-artesunate structure (PDB:6FGC). All figures representing protein structures were generated with PyMOL (Schrodinger LLC), Chimera (Pettersen et al., 2004) and ChimeraX (Goddard et al., 2018).

#### Enzymatic Activity Assay

Pyridoxal kinase activity (WT and variants) were measured following a previously described procedure (Kwok and Churchich, 1979). Briefly, the assay was conducted in 10 mM Hepes buffer (pH 7.3) at 37°C with 100 mM KCl, 1 mM MgCl_2_, 1 mM Mg-ATP and 50 μg/mL BSA (bovine serum albumin). The pyridoxal kinase concentration was 20 μg/ml (0.6 μM), and the substrate pyridoxal was added in a range from 10 μM up to 600 μM. The activity was measured following the increase in absorbance at 388 nm due to PLP formation (extinction coefficient of 4900 M^-1^cm^-1^) in a CLARIOstar (BMG LABTECH) microplate reader. All experiments were carried out in triplicates. K_M_ and k_cat_ values were calculated by a Lineweaver-Burk plot (Lineweaver and Burk, 1934) with the program Prism (GraphPad Software). For statistical significance of the enzymatic assays, initially, the normality distribution of the data was determined by a D’Agostino & Pearson normality test. After passing the normality test, the statistical significance was determined by the paired *t*-test. For all statistical tests, the p values correspond to *p<0.05; **p<0.01; ***p<0.001; ****p<0.0001; ns is not significant. Statistical analyses were performed by using values from four independent experiments.

To derive the K_i_ values, the assay was performed under the same conditions using pyridoxal at concentrations of 50 μM and 150 μM. Using both pyridoxal concentrations the assays were performed with a 2-fold serial dilution of artesunate and artemisinin, starting at concentrations of 2.5 mM and 0.156 mM, respectively. K_i_ values for the inhibitors artesunate and artemisinin were estimated by a Dixon plot (Dixon, 1953), by using a linear regression fit (p< 0.0001) of the inverted velocity values. The K_i_ value corresponds to the intersection between the two lines obtained for each individual pyridoxal concentration. For determining the IC_50_ values, the values of inhibitor concentration were transformed to a logarithmic scale and fitted using a nonlinear regression fit with variable slope. IC_50_ values were calculated as the concentration of inhibitor that gives a velocity half way between the minimal and maximal values of the curve. All curve fitting procedures and statistical analyses were performed using Prism (GraphPad Software).

#### Electrophysiology

Transverse hippocampal slices (350 µm thick) were prepared from adult C57Bl/6J mice. Animals were housed under standard conditions and all procedures were conducted according to the guidelines and with approval of the local government. Whole-cell voltage-clamp recordings were obtained from visualized pyramidal cells of the hippocampal CA1 region in a submerged chamber with perfusion solution containing (in mM) 125 NaCl, 3 KCl, 2.5 CaCl_2_, 1.5 MgCl_2_, 1.25 NaH_2_PO_4_, 25 NaHCO_3_ and 10 D-glucose at 31° C, constantly gassed with 95% O_2_ - 5% CO_2_ (pH 7.4). Miniature inhibitory postsynaptic currents (mIPSCs) were recorded in the presence of the ionotropic glutamate receptor antagonist kynurenic acid (2 mM) and TTX (0.5 µM) using pipettes filled with solution containing (in mM), 130 CsCl, 5 HEPES, 3 MgCl_2_, 5 EGTA, 2 Na_2_ATP, 0.3 Na_3_GTP, 4 NaCl and 5 QX-314 (pH 7.3). mIPSCs were recorded at a holding potential of −70 mV as downward deflections. Current signals were filtered at 2 kHz and sampled at 20 kHz using a Multiclamp 700B amplifier together with a Digidata 1440A interface and the pClamp10 software (Molecular Devices, Sunnyvale, CA). Data were analyzed off-line with Clampfit 10.6 (Molecular Devices) and were expressed as means ± SEM. Statistical comparisons of drug effects were performed with OriginPro 2015G (OriginLab Corporation, MA, USA) using a paired *t*-test. Significance was assumed for p-values < 0.05.

#### Animals for the hippocampal culture

Adult animals (12 weeks old male or female mice) were taken from the mouse strain CD1 (Strain code: 022, Charles River, Sulzfeld, Germany) to isolate the hippocampi. Experiments were authorized by the local veterinary authority and Committee on the Ethics of Animal Experiments (Regierung von Unterfranken).

#### Fluorimetric assay for GAD activity determination

The basis for the determination of the activity of glutamic decarboxylase (GAD) in brain tissue is the ninhydrin reaction. During this reaction, the substrate ninhydrin reacts with the amino-group of γ-aminobutyric acid (GABA) releasing water and forming a Schiff base. Following decarboxylation of the carboxyl group of the amino acid (GABA) and elimination of the amino acid, amino-ninhydrin is formed, which dimerizes with ninhydrin (blue color). The concentration of ninhydrin is proportional to the concentration of the amino acid (Law of Lambert-Beer) and can be measured with a fluorescence spectrophotometer using an excitation wavelength of 375 nm (6 mm slide width) and an emission wavelength of 450 nm (10 mm slide width) (dynode voltage 500 volts) (FluoroMax®-4, HORIBA Scientific, Bangalore, India) (Holdiness et al., 1980; Lowe et al., 1958).

The experimental setup was slightly modified from Holdiness et al., 1980. Hippocampi were transferred into a fresh tube containing sonification buffer (0.5 M KCl, 0.01 M EDTA, 0.5% Triton-X100 in sodium-phosphate buffer, pH 6.4) followed by sonification for 10 s. Protein concentration was measured by a Bradford assay and adjusted to 1 µg/µl. The following conditions were analyzed in triplicates: hippocampi alone, DMSO, 3 µM artemisinin, 10 µM artemisinin and 30 µM artemisinin. For each sample a probe (P) and blank (B) was used with 100 µl homogenized hippocampi and the corresponding DMSO/artemisinin concentrations (3 µM, 10 µM, 30 µM) diluted in sonification buffer. The B fraction was supplemented with 200 µl 10% TCA. The P and B fractions were supplemented with 100 µl substrate buffer (100 mM sodium-L-glutamate buffer in 0.4 M sodium phosphate buffer, pH 6.7, 40 µl 50 mM pyridoxal phosphate 5-phosphate buffer) and incubated for 2 h at 38 °C. Subsequently the reaction of the P fraction was also stopped by adding 200 µl 10% TCA. Samples were centrifuged at 950 x g for 20 min. 200 µl of all probes were incubated with 400 µl ninhydrin (14 mM in 0.5 M sodium carbonate buffer, pH 9.93) at 60°C for 30 min. Afterwards, probes were incubated with 9 ml copper tartrate (1.6 g sodium carbonate, 329 mg tartaric acid, 300 mg copper-(II) sulfate in 1 L *aqua dest*) at 22 °C for 20 min and measured (1:10 diluted) in the fluorescence spectrophotometer. A GABA concentration series of 0, 0.5, 1, 3, 5, 20, 50 µM GABA was used as standard.

### Preparation of primary hippocampal neurons

Hippocampal neurons were prepared at embryonic day 17 (E17) from pregnant female wild type mice. Dissociated cells were grown in neurobasal medium supplemented with 5 ml of L-glutamine (200 mM) and B27 supplement (Life Technologies, A3582801, Germany) with an exchange of 50% medium after 6 days in culture.

### Protein lysate preparation

At day *in vitro* (DIV) 14, hippocampal neurons were incubated for 2 h with different DMSO/artemisinin concentrations (1 µM, 3 µM, 10 µM, 30 µM). Cells were washed and harvested in phosphate-buffered saline pH 7.4 with the help of a cell scraper. After a centrifugation step, the pellet was resuspended in 100 µl of brain homogenate buffer (20 mM Hepes, 100 mM KCH_3_COOH, 40 mM KCl, 5 mM EGTA, 5 mM MgCl_2_, 5 mM DTT, 1 mM PMSF, 1% Triton X, protease inhibitor Roche complete, pH 7.2) and sonicated at low power for 5 s. Protein concentration was determined with the Bradford assay. 10 µg per condition were used for Western Blot analysis.

### Western Blot

For SDS-PAGE, 11% polyacrylamide gels were freshly prepared, followed by Western blot on nitrocellulose membranes (GE Healthcare, Little Chalfont, UK). Membranes were blocked for 1 h with 5% BSA in TBS-T (TBS with 1% Tween 20). Primary antibodies were incubated overnight at 4°C. GAD and PDXK proteins were detected with the GAD67/65 specific antibody (ab11070 1:1,000, abcam, Berlin, Germany) and PDXK specific antibody (NBP1-88283, 1:1000, novusBio, Wiesbaden, Germany). β-Actin (GTX26276, 1:5,000, GeneTex/Biozol, Irvine, CA, USA) served as loading control. Signals were detected using the ECL plus system (GE Healthcare, Little Chalfont, UK).

### Data analysis of Western blots

The image quantification was performed using ImageJ (1.51)/Fiji^2^ (Schindelin et al., 2012, 2015; Schneider et al., 2012). The data were analyzed using Student’s *t*-test (analysis of variance) and values below _*_*p* < 0.05 were considered significant, _**_*p* < 0.01, _***_*p* < 0.001. The values are displayed as mean ± standard error of the mean (±SEM) or as otherwise noted.

### Immuncytochemical staining

DIV14 primary hippocampal neurons were incubated for 2 h with corresponding DMSO/artemisinin concentrations (1 µM, 3 µM, 10 µM). Neurons were fixed in 4% paraformaldehyde in PBS for 15 min. After washing twice with PBS, 50 mM NH_4_Cl was added for 10 min. Blocking with 5% goat serum in PBS (permeabilized with 0.2% Triton X-100) for 30 min at 22°C followed. Primary antibodies were incubated for 1 h in blocking solution without Triton X-100. GAD, gephyrin and PDXK proteins were detected with the GAD67/65 specific antibody (ab11070 1:500, abcam, Berlin, Germany), gephyrin specific antibody (147111, 1:500, Synaptic Systems, Göttingen, Germany) and PDXK specific antibody (NBP1-88283, 1:150, novusBio, Wiesbaden, Germany), respectively. Secondary antibodies gαmCy3, gαrCy5 (1:500; Dianova, Hamburg, Germany) and ActinGreenTM (R37110, Thermo Fisher Scientific, Irvine, CA, USA) were applied for 1 h. Cells were stained with 4’,6-diamino-2-phenylindole (DAPI) and slides were mounted with Mowiol.

## QUANTIFICATION AND STATISTICAL ANALYSIS

The programs and software used for quantification and statistical analysis are mentioned in the methods details section in detail. Statistical analyses in this manuscript are described in the experimental methods section and also in the corresponding figure legends.

## DATA AVAILABILITY

The coordinates of mPDXK-apo, mPDXK-ATPγS and mPDXK-ATPγS-artesunate structures have been deposited in the Protein Data Bank (PDB) with accession codes 6YJZ, 6YK0 and 6YK1, respectively.

## SOURCE TABLE

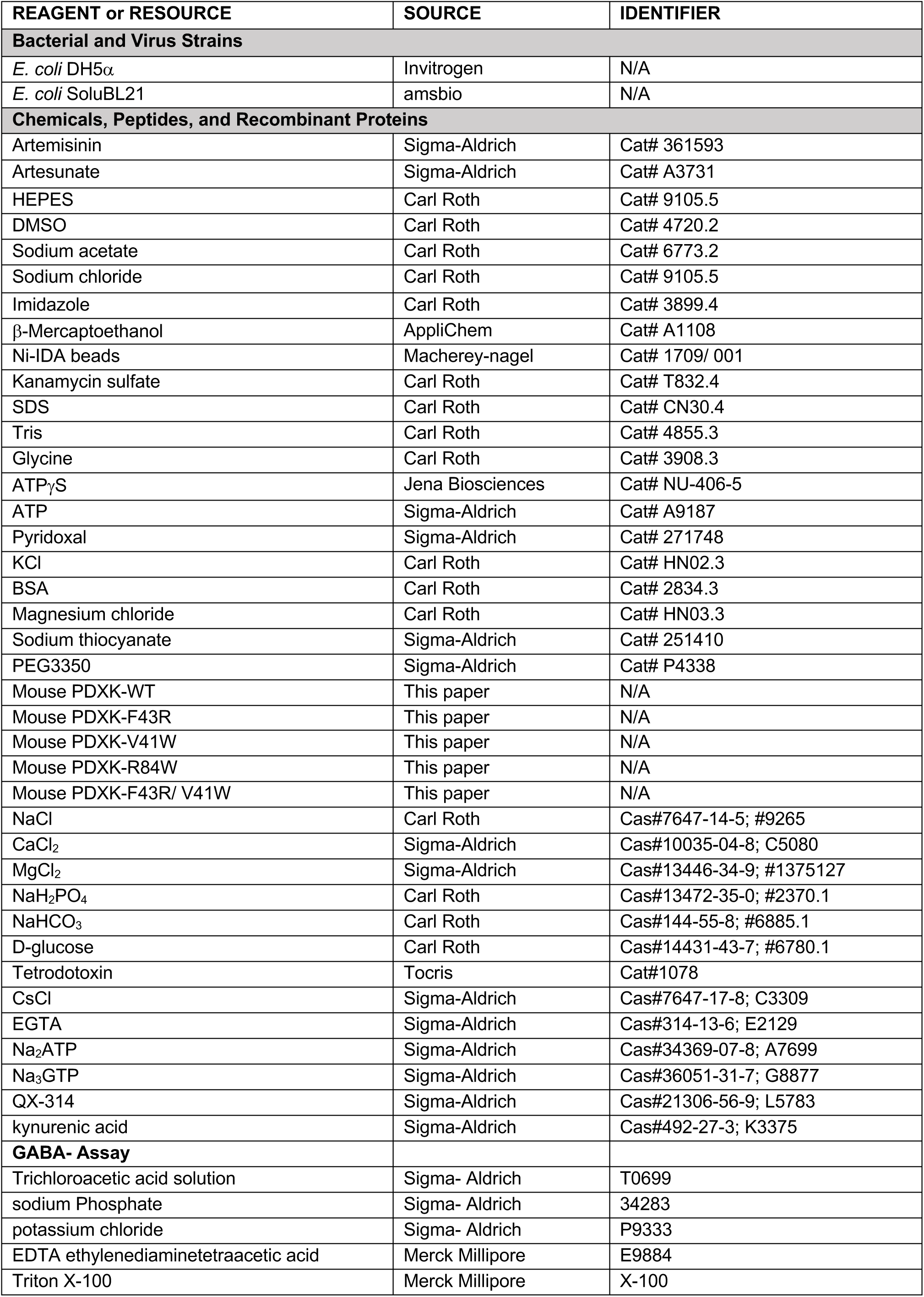
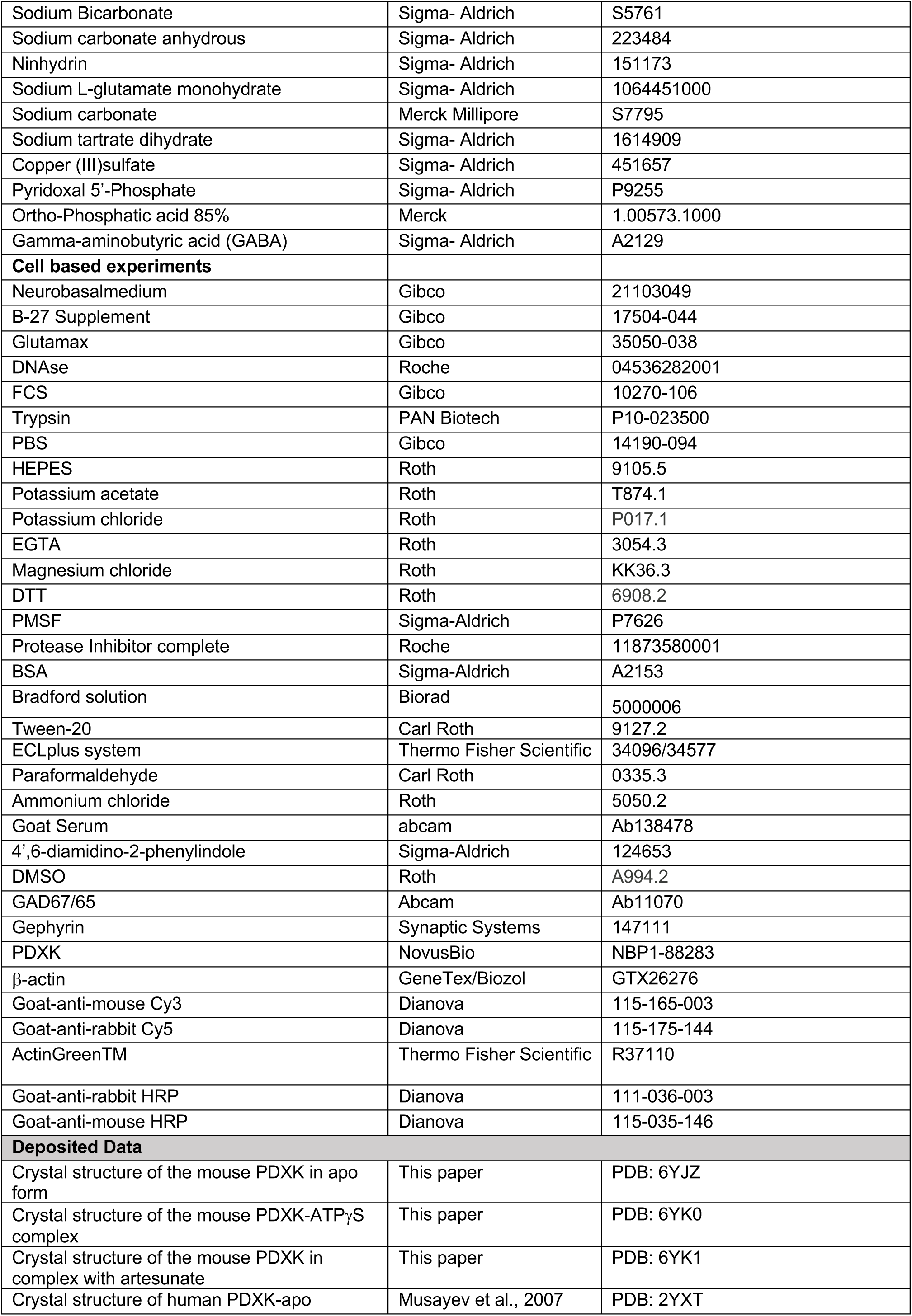
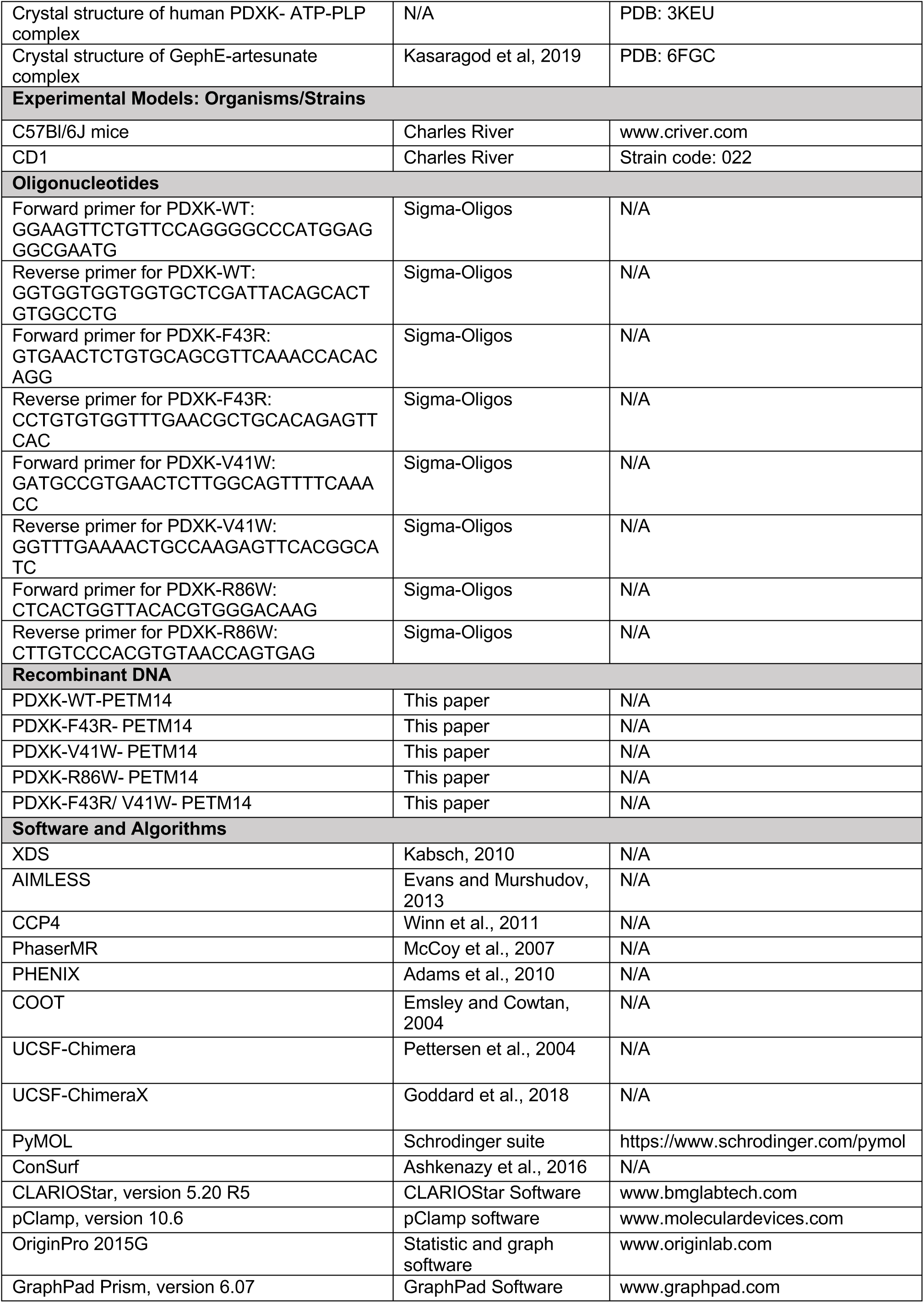

## Figure supplements

**Figure 1 – figure supplement 1.**
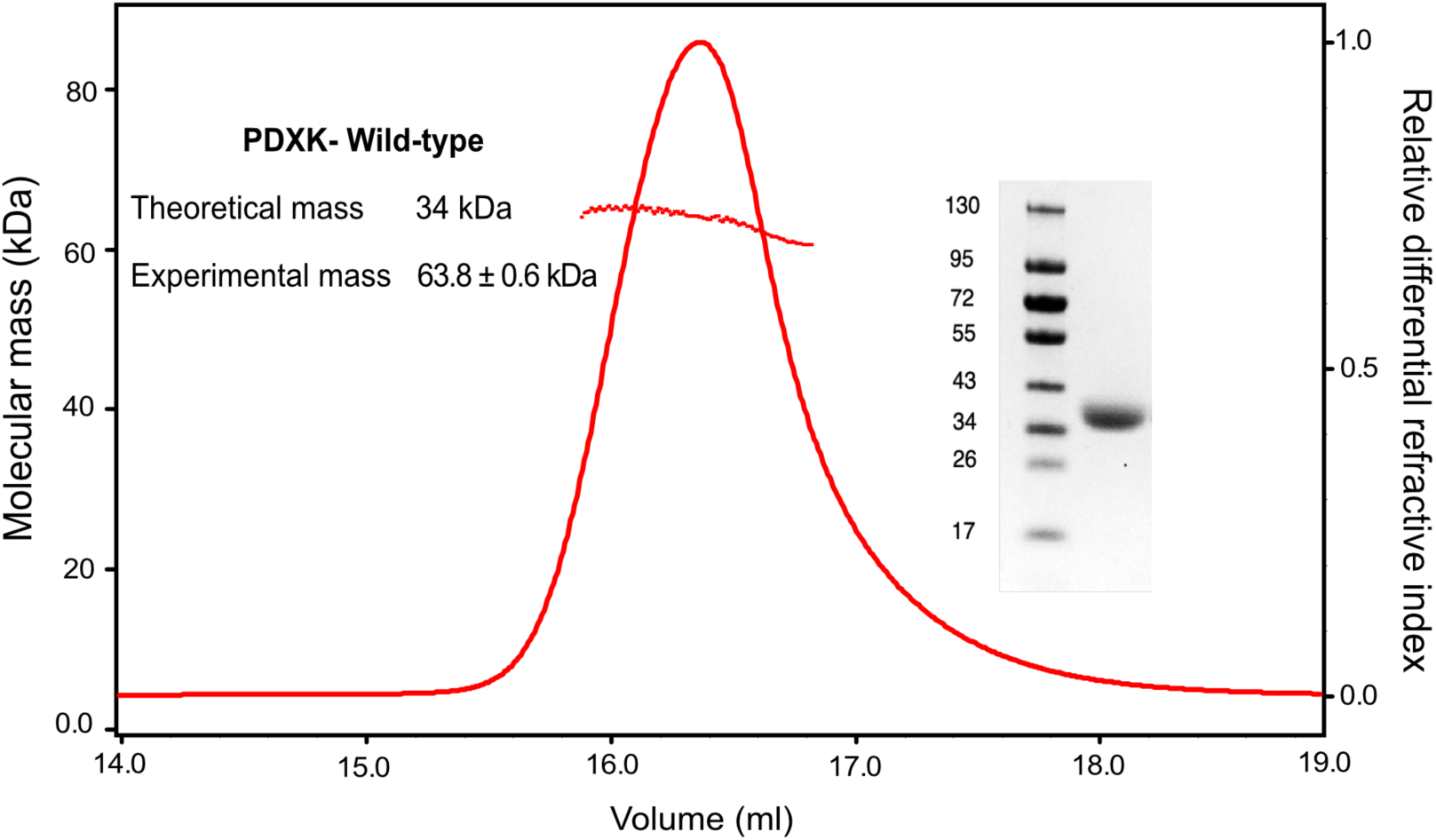
MALS measurements of WT PDXK. Multi angle laser light scattering coupled to size exclusion chromatography (SEC-MALS) of recombinantly expressed and purified WT-PDXK along with an SDS-PAGE analysis of the purified protein (inset). Normalized differential refractive index is represented along with the measured molecular mass under the curve. The SEC-MALS experiment demonstrates that the protein is dimeric in solution.

**Figure 1 – figure supplement 2.**
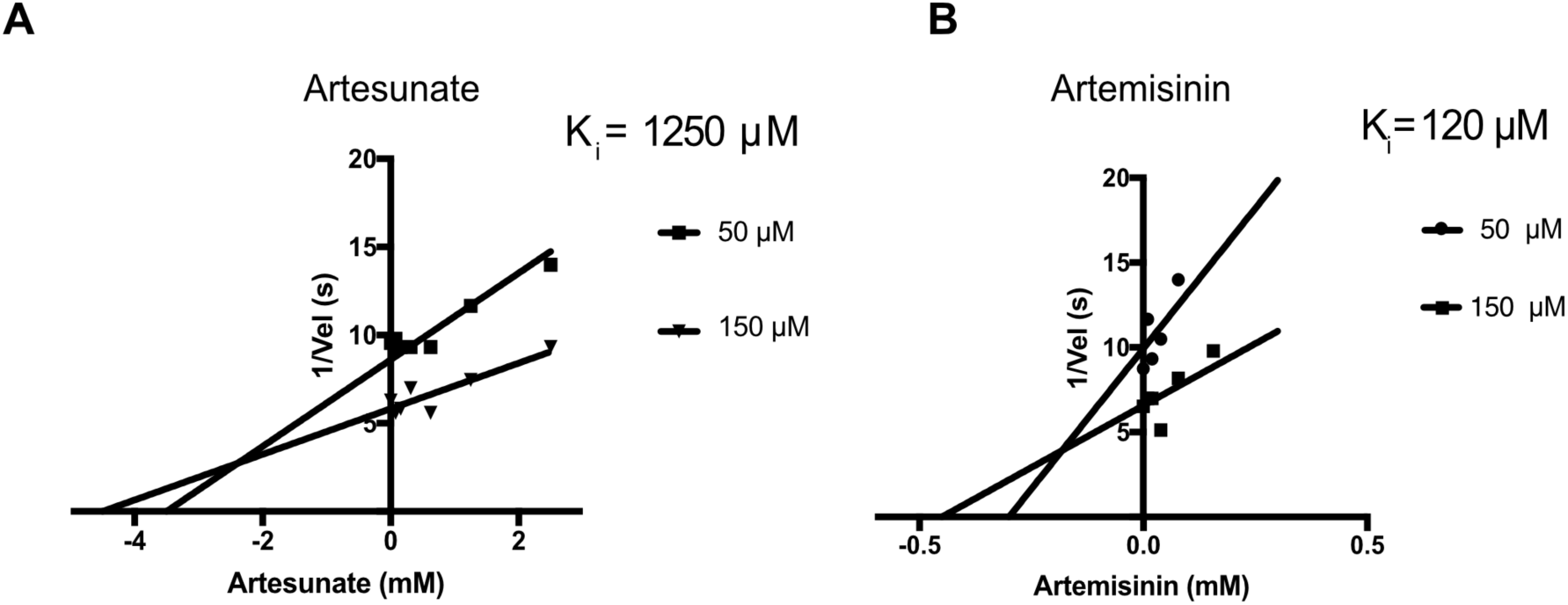
Inhibition analysis. **(A-B)** Dixon plots for the inhibition of PDXK by artesunate **(A)** and artemisinin **(B)**. The succinate derivative of artemisinin displays a K_i_ of 1250 μM in contrast to the more potent artemisinin with a K_i_ of 120 μM.

**Figure 2 – figure supplement 1.**
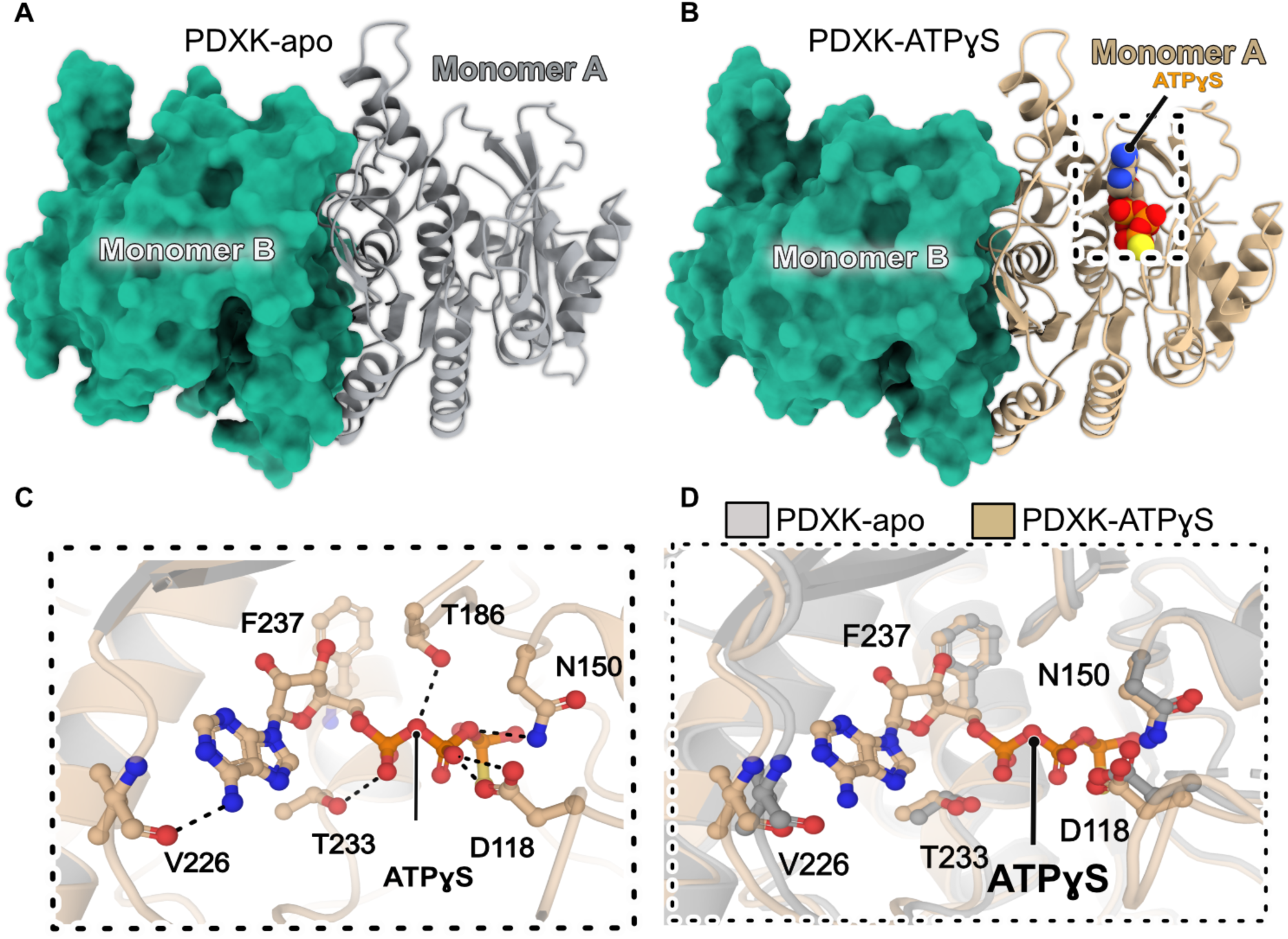
Structural comparison of apo-PDXK and the PDXK-ATPγS complex. **(A-B)** Overall architecture of apo-PDXK (A) and the PDXK-ATPγS (B) complex. One protomer is shown in cartoon and the second in surface representation. ATPγS is displayed in CPK representation. **(C)** Enlarged view of the ATPγS binding pocket. The ligand and residues crucial for binding are shown in ball and stick representation and the protein backbone as a cartoon. **(D)** Comparative analyses of the PDXK-apo and PDXK-ATPγS structures. One monomer of PDXK-apo and PDXK-ATPγS are displayed in gray and brown cartoon representation, respectively, while the other monomer is displayed in surface representation in green.

**Figure 2 – figure supplement 2.**
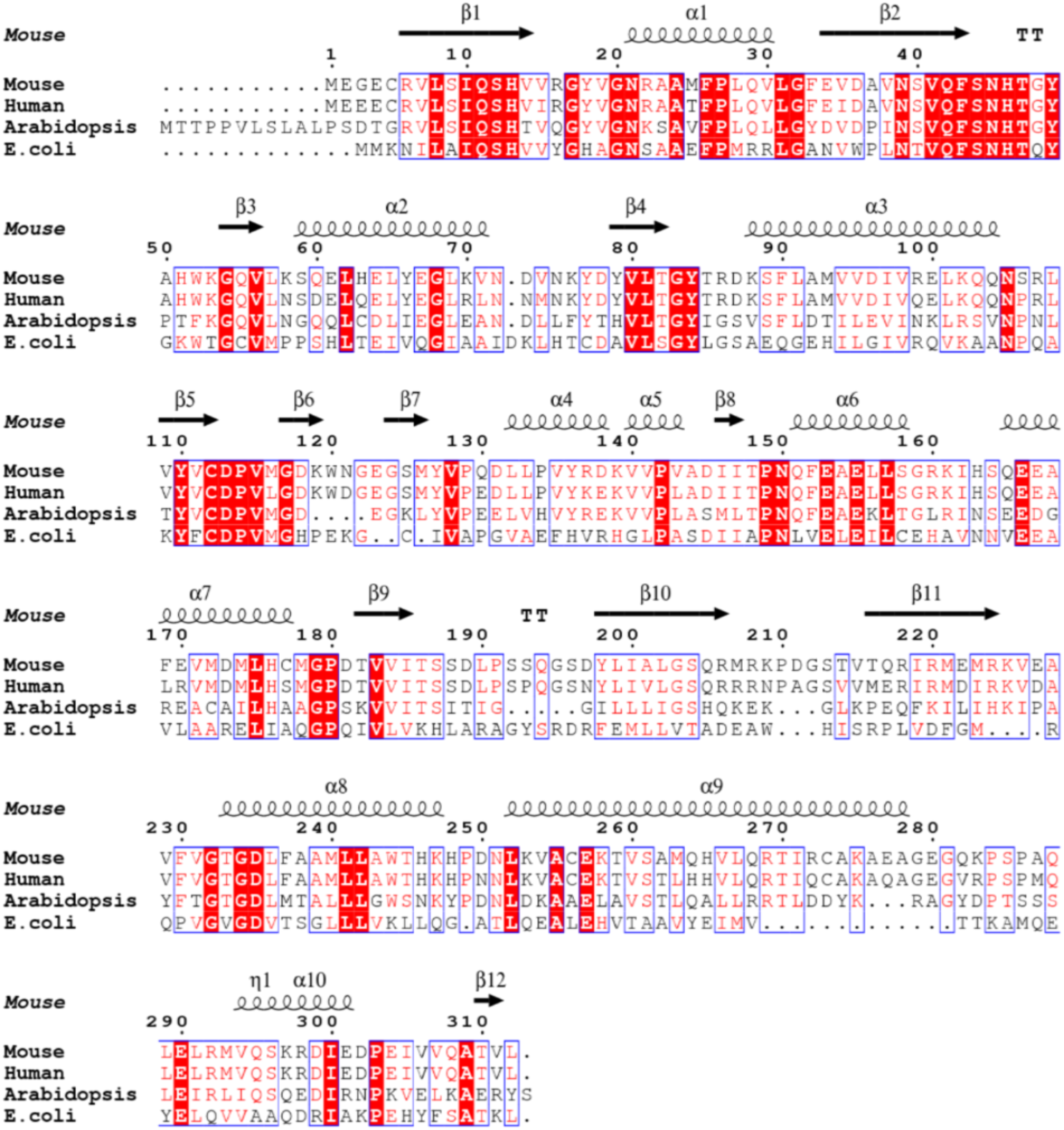
Multiple sequence alignment. Multiple sequence alignment of PDXK enzymes from diverse sources obtained with Clustal omega (Sievers et al., 2011) and represented by using the ESPript server (Robert and Gouet, 2014).

**Figure 2 – figure supplement 3.**
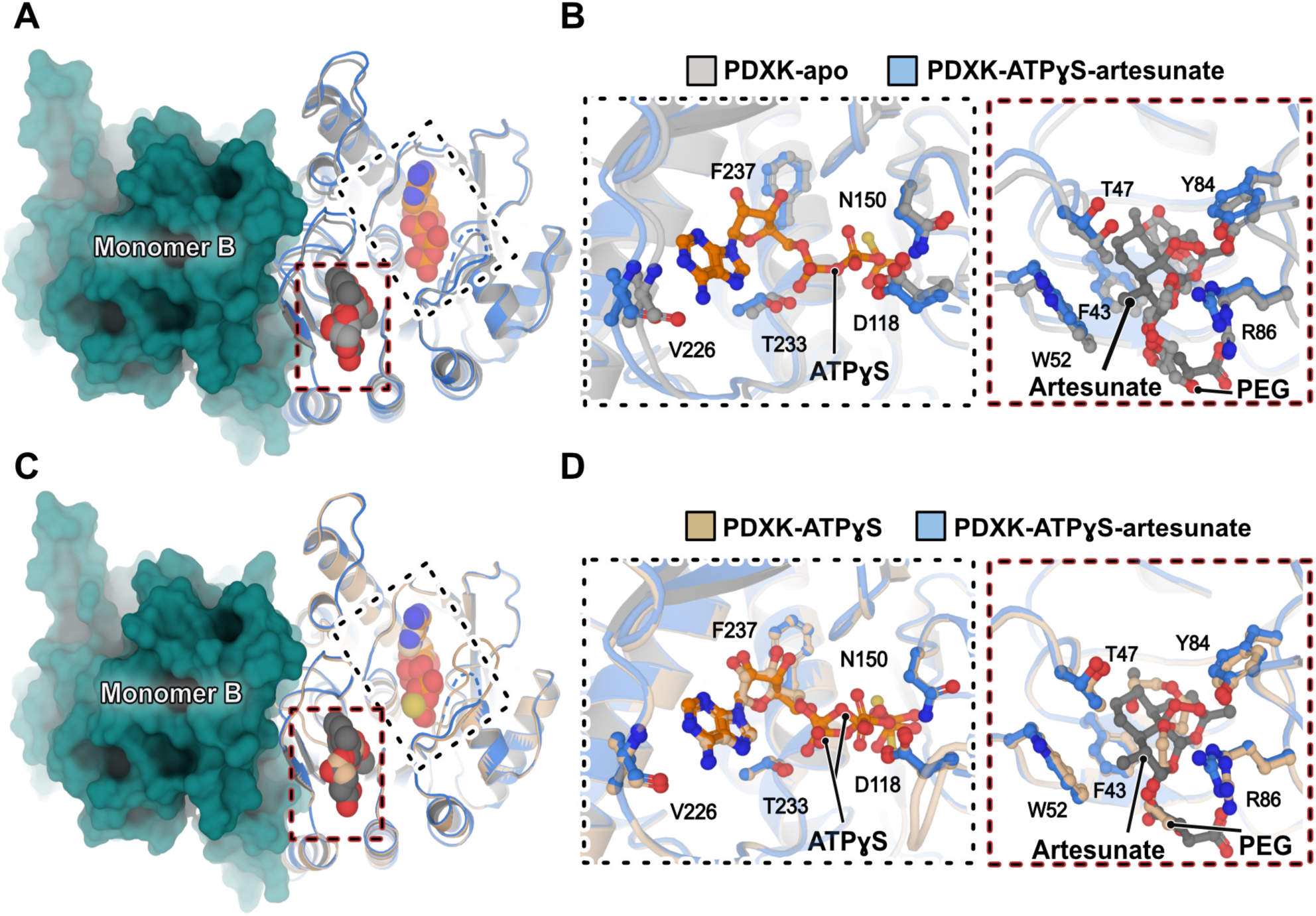
Structural comparison of the apo, the PDXK-ATPγS and the PDXK-ATPγS-artesunate structures. **(A-B)** Comparison of the PDXK-ATPγS-artesunate structure with the apo structure. The overall architecture is shown in **(A)** and enlarged views of the ligand binding pocket are shown in **(B). (C-D)** Comparison of the PDXK-ATPγS-artesunate structure with the PDXK-ATPγS complex. The overall architecture is shown in **(C)** and enlarged views of the ligand binding pocket are shown in **(D)**. In panels **B** and **D**, bound ligands and residues, which are crucial for binding, are shown in ball and stick representation and the protein backbone in cartoon representation. Please note that in the apo and in the binary PDXK-ATPγS structure, the artemisinin binding pocket is occupied by polyethylene glycol, a component of the crystallization solution.

**Figure 2 – figure supplement 4.**
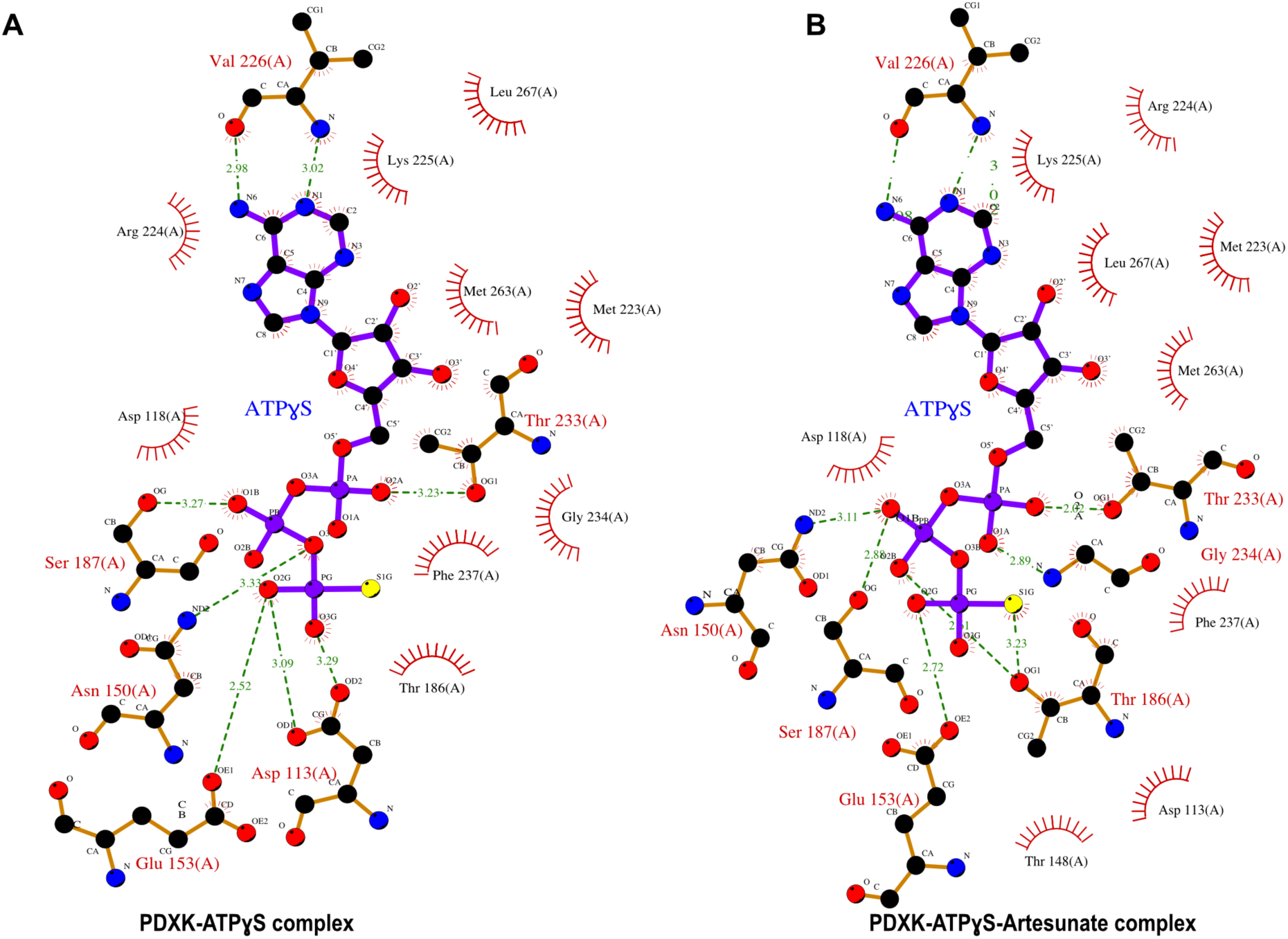
Comparison of ATP binding pockets. **(A-B)** LigPlot 2D-representation of ATPγS from the binary PDXK-ATPγS **(A)** and ternary PDXK-ATPγS-artesunate **(B)** complexes derived from an analysis with the ProFunc server (Laskowski et al., 2005). Please note that residues mediating the binding of ATPγS are almost identical in both structures as is the conformation of the bound ATPγS (not shown).

**Figure 2 – figure supplement 5.**
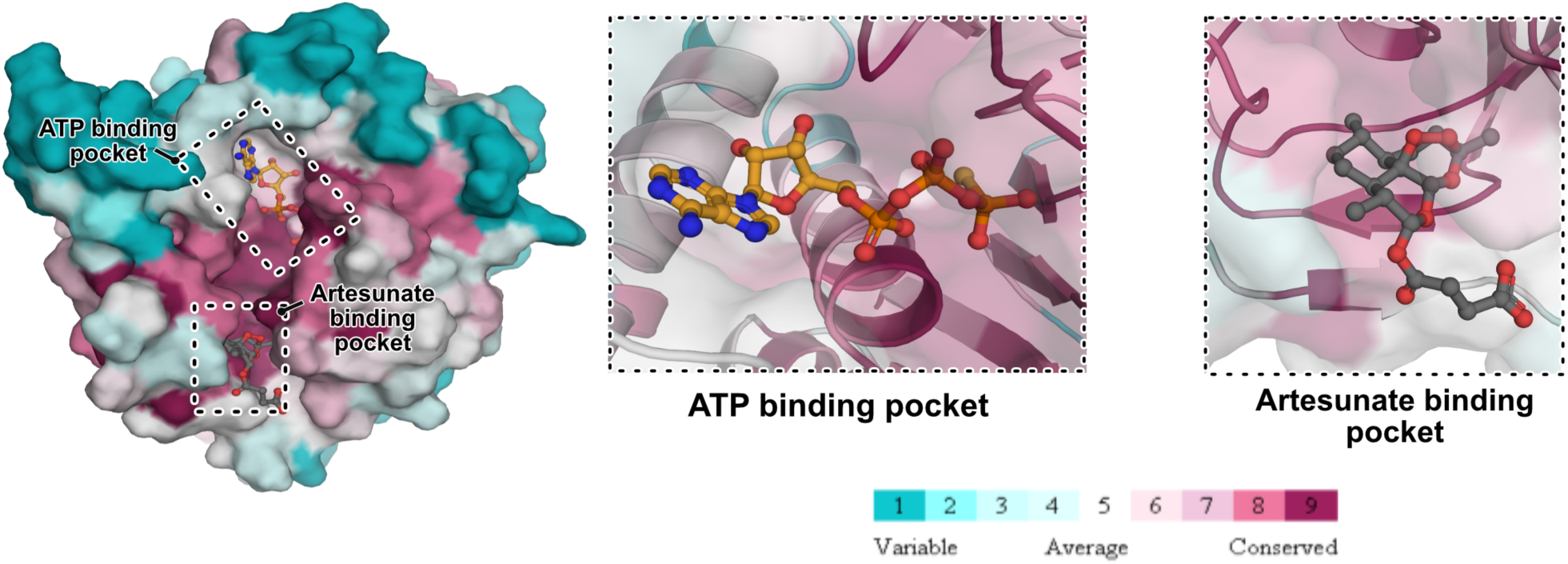
Conservation analysis. PDXK sequence conservation analyzed with the ConSurf server (Ashkenazy et al., 2016). Overall architecture of a PDXK monomer colored according to the accompanying conservation scores. Enlarged views of the ATP and the artesunate binding pockets clearly show that both binding pockets are evolutionarily highly conserved.

**Figure 3 – figure supplement 1.**
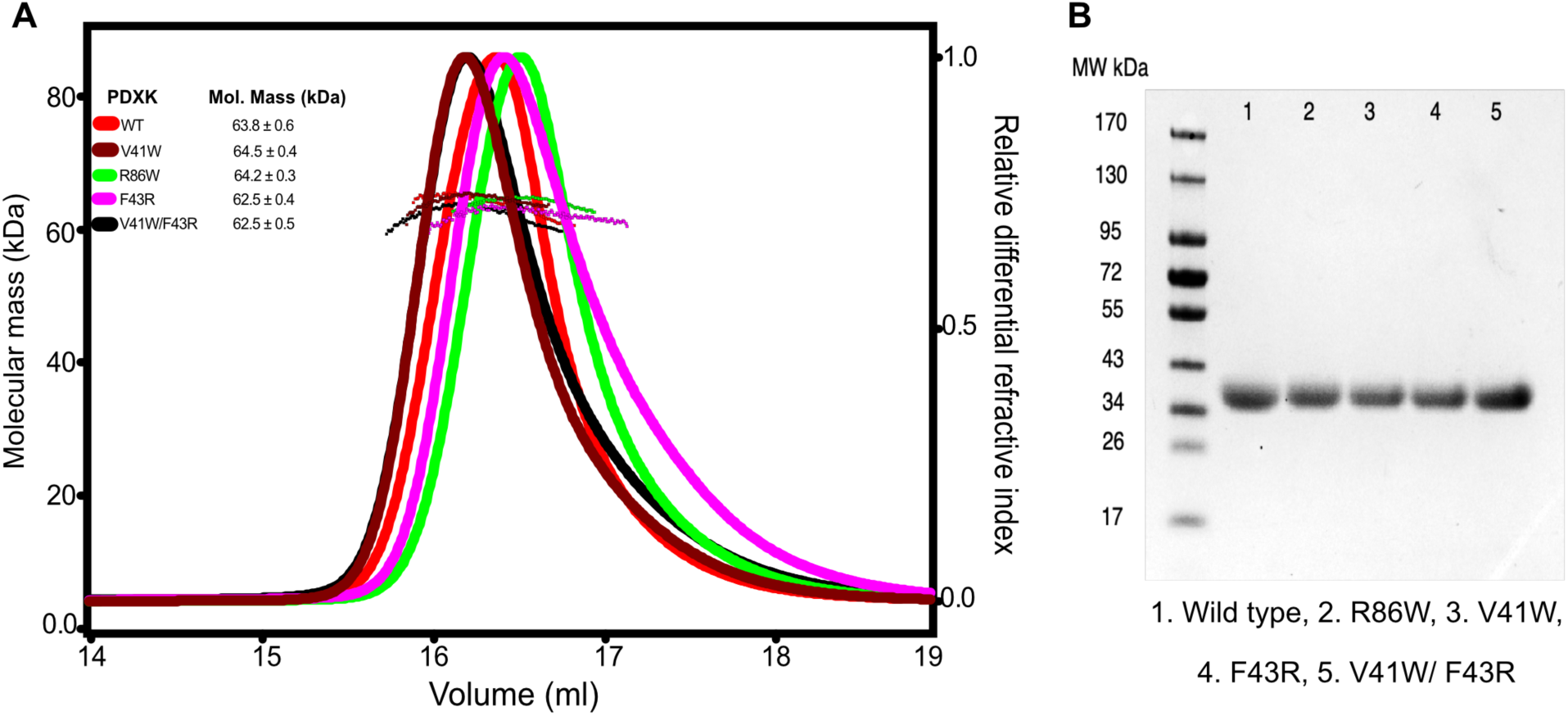
Biophysical and biochemical analysis of PDXK variants. **(A)** SEC-MALS analyses of the wildtype and PDXK mutants. The data confirm that, like the wild-type protein, all mutants dimerize. Please note that due to non-synchronized injections the elution volumes of the different samples are slightly offset. **(B)** SDS-PAGE analysis of purified PDXK variants and the wild-type.

**Figure 3 – figure supplement 2.**
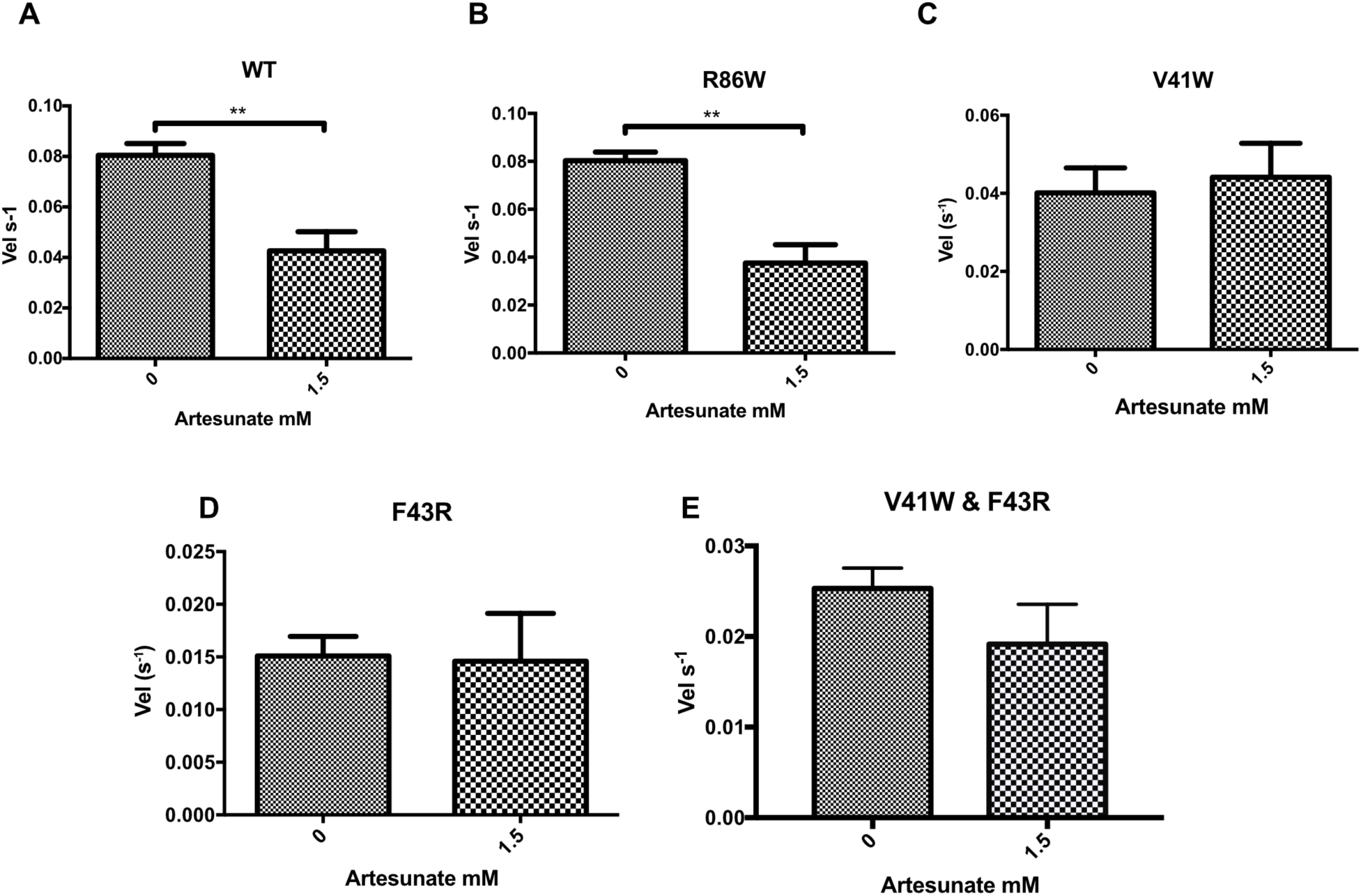
Inhibition analysis of PDXK variants. **(A-E)** Bar diagrams of the turnover rates of PDXK variants in the absence and presence of artesunate (1.5 mM). Please note that R86W **(B)** behaves similar as the WT **(A)** with artesunate retaining its inhibition potency whereas the variants V41W **(C)**, F43R **(D)** and the double mutant V41W/F43R **(E)** completely abolish artesunate binding. Data are presented as mean ± SEM (p values are: *p<0.05; **p<0.01; ***p<0.001; ****p<0.0001, Paired *t* test).

**Figure 5 – figure supplement 1.**
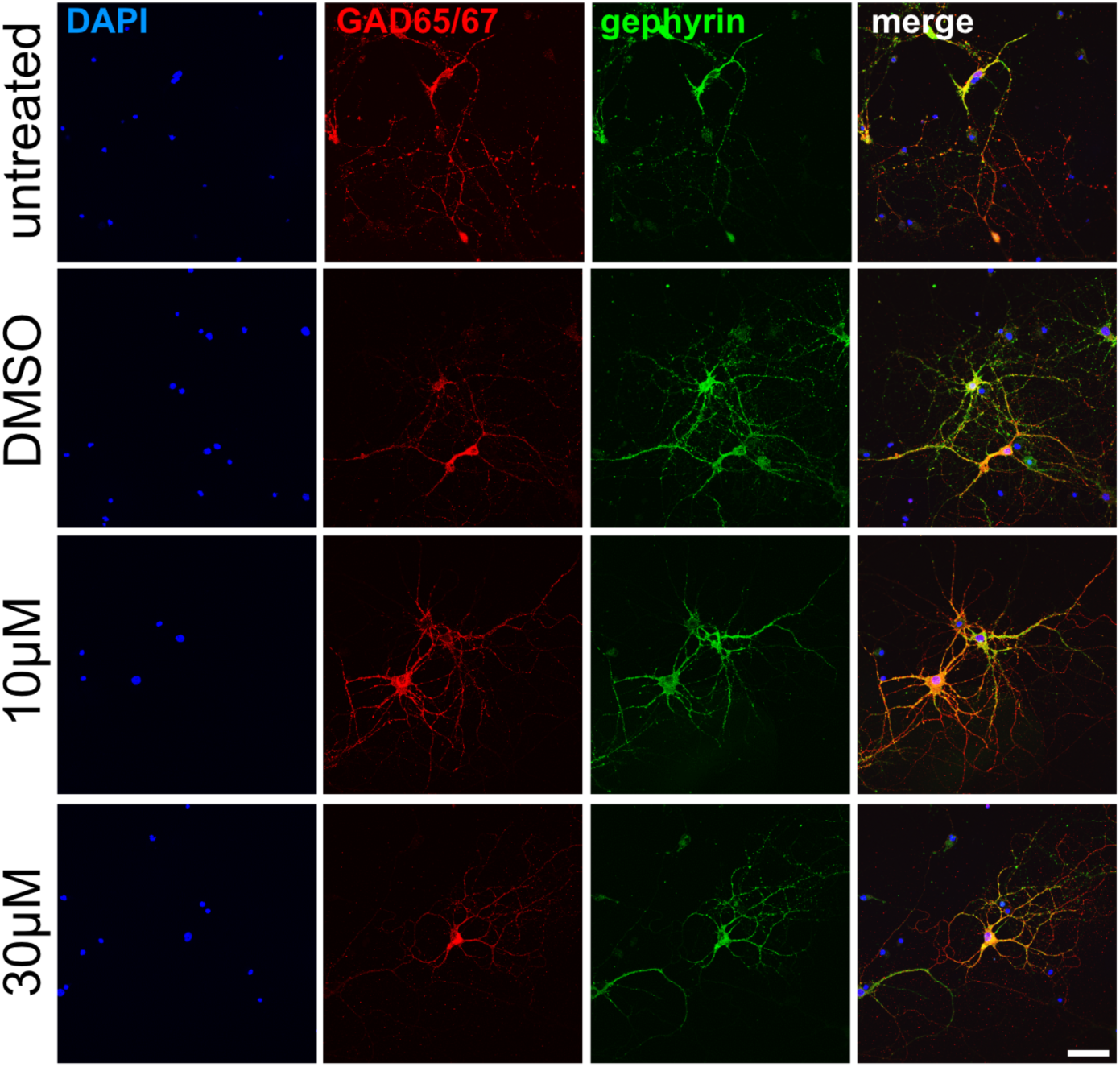
Immunocytochemical staining of GAD and gephyrin. Immunocytochemical staining of hippocampal neurons were performed at DIV14. Untreated cells, DMSO treated controls, and artemisinin (10 µM, 30 µM) treated cells are shown. Cells were stained for GAD (red), gephyrin (green), and with DAPI (blue) to label the nucleus. The merge of all channels is presented in the right panels (yellow overlay of GAD and gephyrin). Scale bar corresponds to 50 µm.

**Figure 5 – figure supplement 2.**
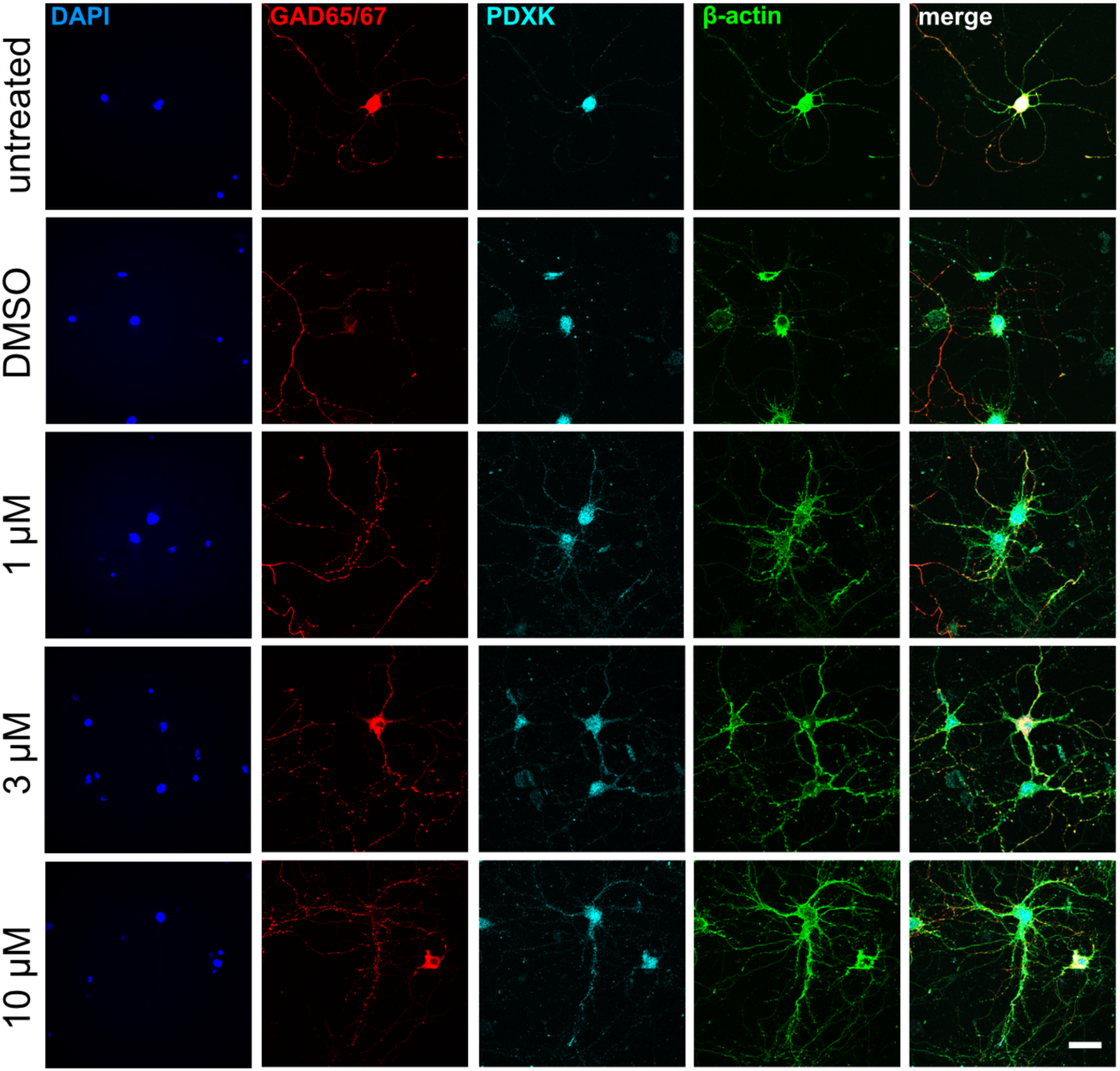
Immunocytochemical staining of PDXK and GAD. Hippocampal neurons at DIV14 were stained for GAD (red), PDXK (cyan), β-actin (green), and DAPI (blue) to label the nucleus. The merge of all channels is presented in the right panels. Untreated cells, DMSO treated controls, and artemisinin (1 µM, 3 µM, 10 µM) treated cells are shown. Note, no change in GAD and PDXK intensity was observed between the different conditions. Scale bar corresponds to 50 µm.

**Figure 5 – figure supplement 3.**
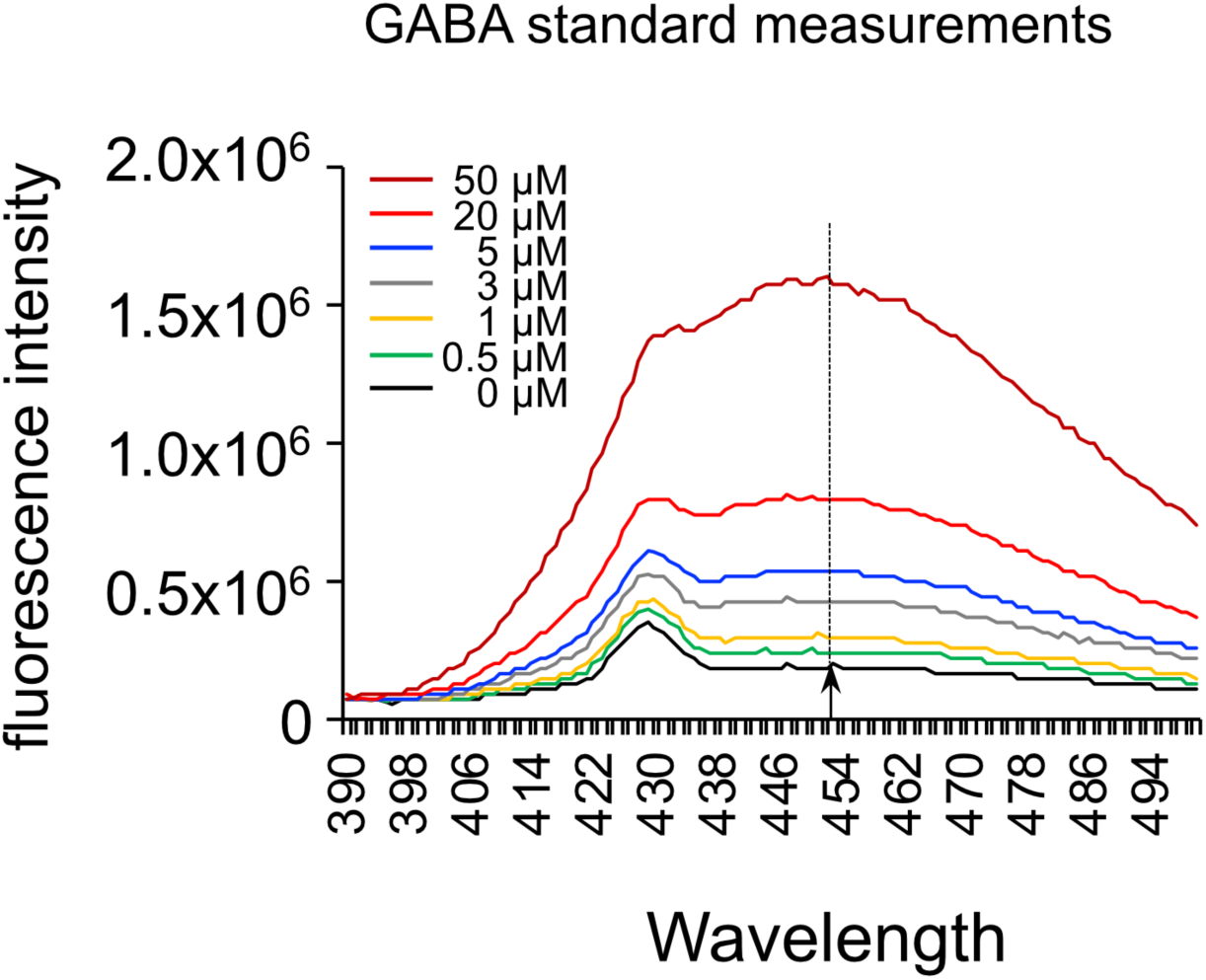
GABA standard measurements. A concentration series of GABA was established (0, 0.5, 1, 3, 5, 20, 50 µM) by measuring the fluorescence spectrum from 390 to 500 nm. GAD activity was quantified based on the emission at 450 nm (marked by an arrow and dotted line). These standard values were used to measure the amount of GABA synthesized by GAD as represented in **Figure 5D**.

